# svclassify: a method to establish benchmark structural variant calls

**DOI:** 10.1101/019372

**Authors:** Hemang Parikh, Marghoob Mohiyuddin, Hugo Y. K. Lam, Hariharan Iyer, Desu Chen, Mark Pratt, Gabor Bartha, Noah Spies, Wolfgang Losert, Justin M. Zook, Marc Salit

**Affiliations:** Genome-Scale Measurements Group, National Institute of Standards and Technology, Gaithersburg, MD 20899, USA.; Dakota Consulting Inc., 1110 Bonifant Street, Suite 310, Silver Spring, MD 20910, USA.; Bina Technologies, Roche Sequencing, Redwood City, CA 94065, USA.; Statistical Engineering Division, National Institute of Standards and Technology, Gaithersburg, MD 20899, USA.; Institute for Research in Electronics and Applied Physics, University of Maryland, College Park, MD 20742, USA.; Personalis Inc., 1350 Willow Road, Suite 202, Menlo Park, CA 94025, USA.; Department of Pathology, Stanford University, Stanford, CA, USA.; Bioengineering Department, Stanford University, Stanford, CA, USA.

## Abstract

**Background:** The human genome contains variants ranging in size from small single nucleotide polymorphisms (SNPs) to large structural variants (SVs). High-quality benchmark small variant calls for the pilot National Institute of Standards and Technology (NIST) Reference Material (NA12878) have been developed by the Genome in a Bottle Consortium, but no similar high-quality benchmark SV calls exist for this genome. Since SV callers output highly discordant results, we developed methods to combine multiple forms of evidence from multiple sequencing technologies to classify candidate SVs into likely true or false positives. Our method (svclassify) calculates annotations from one or more aligned bam files from many high-throughput sequencing technologies, and then builds a one-class model using these annotations to classify candidate SVs as likely true or false positives.

**Results:** We first used pedigree analysis to develop a set of high-confidence breakpoint-resolved large deletions. We then used svclassify to cluster and classify these deletions as well as a set of high-confidence deletions from the 1000 Genomes Project and a set of breakpoint-resolved complex insertions from Spiral Genetics. We find that likely SVs cluster separately from likely non-SVs based on our annotations, and that the SVs cluster into different types of deletions. We then developed a supervised one-class classification method that uses a training set of random non-SV regions to determine whether candidate SVs have abnormal annotations different from most of the genome. To test this classification method, we use our pedigree-based breakpoint-resolved SVs, SVs validated by the 1000 Genomes Project, and assembly-based breakpoint-resolved insertions, along with semi-automated visualization using svviz.

**Conclusions:** We find that candidate SVs with high scores from multiple technologies have high concordance with PCR validation and an orthogonal consensus method MetaSV (99.7% concordant), and candidate SVs with low scores are questionable. We distribute a set of 2676 high-confidence deletions and 68 high-confidence insertions with high svclassify scores from these call sets for benchmarking SV callers. We expect these methods to be particularly useful for establishing high-confidence SV calls for benchmark samples that have been characterized by multiple technologies.

## Background

The human genome contains variants ranging in size from small single nucleotide polymorphisms (SNPs) to large structural variants (SVs). SVs include variations such as novel sequence insertions, deletions, inversions, mobile-element insertions, tandem duplications, interspersed duplications and translocations. In general, SVs include deletions and insertions larger than 50 base pairs (bps), while smaller insertions or deletions are referred to as indels, though the threshold of 50 bps is somewhat arbitrary and based on the fact that different bioinformatics methods are usually used to detect SVs vs. small indels and SNPs. SVs have long been implicated in phenotypic diversity and human diseases [1]; however, identifying all SVs in a whole genome with high-confidence has proven elusive. Recent advances in next-generation sequencing (NGS) technologies have facilitated the analysis of SVs in unprecedented detail, but these methods tend to give highly non-overlapping results [2]. In this work, we calculate “annotations” from features in the reads in and around candidate SVs, and we then develop methods to evaluate candidate SVs based on evidence from multiple NGS technologies.

NGS offers unprecedented capacity to detect many types of SVs on a genome-wide scale. Many bioinformatics algorithms are available for detecting SVs using NGS including depth of coverage (DOC), paired-end mapping (PEM), split-read, junction-mapping, and assembly-based methods [2]. DOC approaches identify regions with abnormally high or low coverage as potential copy number variants. Hence, DOC methods are limited to detecting only deletions and duplications but not other types of SVs, and they have more power to detect larger events and deletions. PEM methods evaluate the span and orientation of paired-end reads. Read pairs map farther apart around deletions and closer around insertions, and orientation inconsistencies indicate potential inversions or tandem duplications. Split reads are used to identify SVs by identifying reads whose alignments to the reference genome are split in two parts and contain the SV breakpoint. Junction-mapping methods map poorly mapped, soft-clipped or unmapped reads to junction sequences around known SV breakpoints to identify SVs. Assembly-based methods first perform a *de novo* assembly, and then the assembled genome is compared to the reference genome to identify all types of SVs. By combining various approaches to detect SVs, it is possible to overcome the limitations of individual approaches in terms of the types and sizes of SVs that they are able to detect, but still difficult to determine which are true [3, 4, 5].

Numerous methods have been developed to find candidate SVs using NGS, but clinical adoption of human genome sequencing requires methods with known accuracy. The Genome in a Bottle Consortium (GIAB) is developing well-characterized whole-genome reference materials for assessing variant-call accuracy and understanding biases. Recently GIAB released high-confidence SNP, indel, and homozygous reference genotypes for Coriell DNA sample NA12878, which is also National Institute of Standards and Technology (NIST) Reference Material 8398 available at https://www-s.nist.gov/srmors/view_detail.cfm?srm=8398 [6]. In this work, we developed methods to integrate evidence of SVs in mapped sequencing reads from multiple sequencing technologies. We used unsupervised machine learning to determine the characteristics of the different SV types, and we used One Class Classification to classify candidate SVs as likely true positives, false positives, or ambiguous. Using these methods, we classified three independently established “validated” call sets containing large deletions or insertions.

Our classification methods use the machine learning technique One Class Classification (OCC) [7, 8]. In contrast to the more common two-class models that have two training sets (e.g., positives and negatives), one-class methods have only a single training set and try to identify sites unlike the training set. In our OCC methods, the algorithm tries to identify a region, R, of the annotation space that contains a specified, large proportion (e.g. 95% or 99%) of the non-SVs. Sites that have annotations falling outside R are classified as SVs. In essence, these are outliers relative to the non-SVs. For selecting R, only a representative set of non-SVs is required for the training. In our model, we use random genomic coordinates as our one class because random coordinates are unlikely to be near true SV breakpoints. For our one-class model, we only include annotations that are likely to indicate a SV if they differ from random coordinates for a defined set of parameters (e.g., read clipping, pair distance, and coverage). We do not include annotations like mapping quality that may not always distinguish SVs from non-SVs because atypical values may also indicate random regions of the genome that are difficult to sequence. We do not use a two class machine learning model because our potential training SV call sets are primarily easier-to-detect mid-size deletions and insertions and are not representative of all types of deletions, insertions, or other SV types, which is an important assumption of two-class models. Therefore, a two class model trying to differentiate our SV sets from random genomic coordinates can do a very good job separating these two sets, but the model is likely to misclassify other candidate SVs not in the “Validated/assembled” call sets (e.g., duplications, deletions in difficult parts of the genome, etc.). Because our one-class model does not rely on biased “Validated/assembled” call sets, we expect our one-class model to be more generalizable to other types of SVs by selecting annotations for which atypical values are usually associated with SVs.

Our methods, which classify based on evidence from multiple technologies, are complementary to the recently published MetaSV method [5], which integrates SVs using multiple bioinformatics methods, and the Parliament method [9], which generates candidate SVs using multiple technologies and bioinformatics methods, and then uses a PacBio/Illumina hybrid assembly to determine whether the candidate SVs are likely to be true. Similar to Parliament, in the characterization of the performance of the LUMPY tool [3], the authors developed a high-confidence set that had breakpoints supported by long reads from PacBio or Moleculo. In addition to using svviz [10] to visualize and determine the number of reads supporting the alternate, we also combine the support from multiple sequencing technologies in a robust machine learning model.

## Results and Discussion

To assess the utility of our classification methods, we compiled four whole genome sequencing datasets for Coriell DNA sample NA12878 (Table 1). We used two deletion call sets from Personalis and the 1000 Genomes Project totaling 3082 unique deletions, as well as 70 assembly-based breakpoint-resolved insertions. Moreover, we generated several likely non-SV call sets with different size distributions and sequence contexts (Table 2). We first generated annotations for the candidate SV and non-SV call sets from the four sequencing datasets. We used hierarchical clustering to show that SVs generally cluster separately from non-SVs using these annotations, and that SVs cluster into several different types of deletions. We used one class classification methods to classify calls as high-confidence SVs or uncertain. To confirm the accuracy of the high-confidence SVs, we present PCR validation results for the Personalis deletions as well as a comparison to mother-father-daughter trio consensus-based calls from MetaSV [5].

**Table 1:**
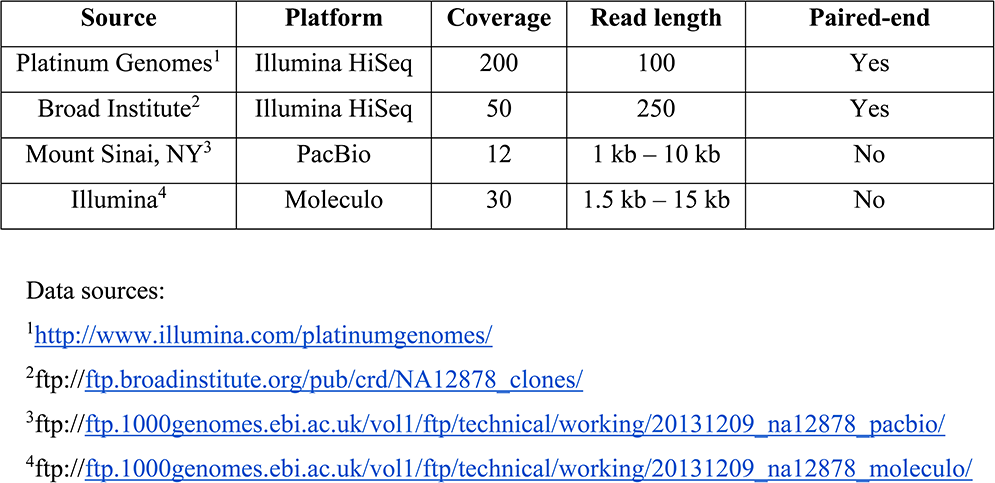
Description of NGS data sets from Coriell DNA sample NA12878.

| Source | Platform | Coverage | Read length | Paired-end |
| --- | --- | --- | --- | --- |
| Platinum Genomes<sup>1</sup> | Illumina HiSeq | 200 | 100 | Yes |
| Broad Institute<sup>2</sup> | Illumina HiSeq | 50 | 250 | Yes |
| Mount Sinai, NY<sup>3</sup> | PacBio | 12 | 1 kb – 10 kb | No |
| Illumina<sup>4</sup> | Moleculo | 30 | 1.5 kb – 15 kb | No |

Data sources:
<sup>1</sup>http://www.illumina.com/platinumgenomes/
<sup>2</sup>ftp://ftp.broadinstitute.org/pub/crd/NA12878_clones/
<sup>3</sup>ftp://ftp.1000genomes.ebi.ac.uk/vol1/ftp/technical/working/20131209_na12878_pacbio/
<sup>4</sup>ftp://ftp.1000genomes.ebi.ac.uk/vol1/ftp/technical/working/20131209_na12878_moleculo/

**Table 2:** Description of SV validated/assembled sets from Coriell DNA sample NA12878.

| Source | # of SVs | # of unique SVs | Size distribution |
| --- | --- | --- | --- |
| Personalis deletions | 2306 | 2292 | 50 to 158654 bps |
| Personalis validated deletions | 39 | 39 | 49 to 9163 bps |
| Personalis non-validated deletions | 5 | 5 | 52 to 7557 bps |
| 1000 Genomes deletions [13] | 2685 | 1825 | 49 to 212899 bps |
| Deduplicated deletions | 3082 | 3082 | 49 to 158654 bps |
| Spiral Genetics insertions | 70 | 70 | 207 to 3865 bps |
| Random regions | 4000 | 4000 | 50 to 997527 bps |
| Random regions (size distribution matching to Personalis) | 2306 | 2306 | 50 to 158654 bps |
| Long interspersed nuclear elements | 497 | 497 | 12 to 6401 bps |
| Long terminal repeat elements | 498 | 498 | 11 to 7511 bps |
| Short interspersed nuclear elements | 496 | 496 | 36 to 335 bps |

### Annotations are generated from multiple technologies for candidate SVs

To assess the evidence for any candidate SV without the need to design primers for validation experiments, we developed svclassify to quantify annotations of aligned reads inside and around each SV (Figure 1). We generated 85 to 180 annotations (Supplementary table 1 to 4) for each of the SV calls as well as likely non-SV regions from four aligned sequence datasets for NA12878 using svclassify. Some of the annotations, such as depth of coverage (Figure 2), could clearly distinguish most Personalis “Gold” deletions from random regions by themselves. Although annotations such as coverage can be used by themselves to classify most Personalis deletions, additional annotations increase confidence that the deletion is real and not an artifact (e.g., low coverage due to extreme GC content). In addition, other annotations are necessary to classify other types of SVs like inversions and insertions that may not have abnormal coverage. Therefore, we developed unsupervised and one-class supervised machine learning models to combine information from many annotations for clustering and classification (Figure 3).

**Figure 1:**
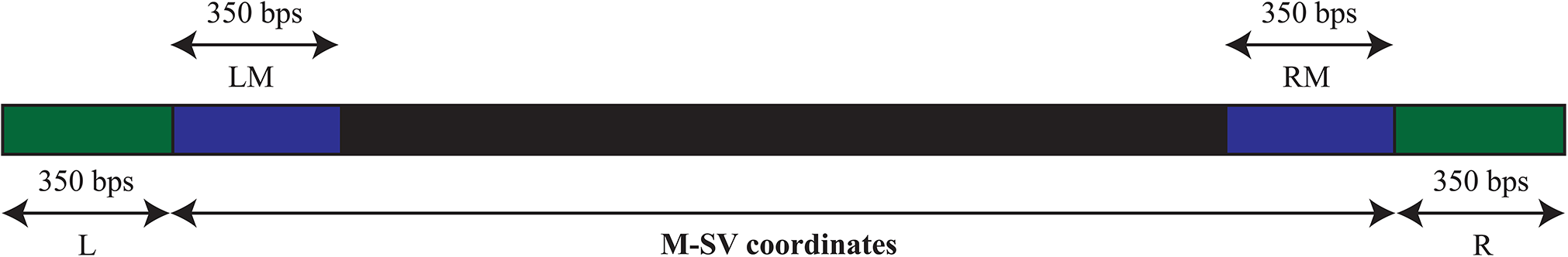
Annotations are generated for each SV for five different regions in and around the SV: Left flanking region (L), Left middle flanking region (LM), Middle regions based on SV coordinates (M), Right middle flanking region (RM), and Right flanking region (R).

**Figure 2:**
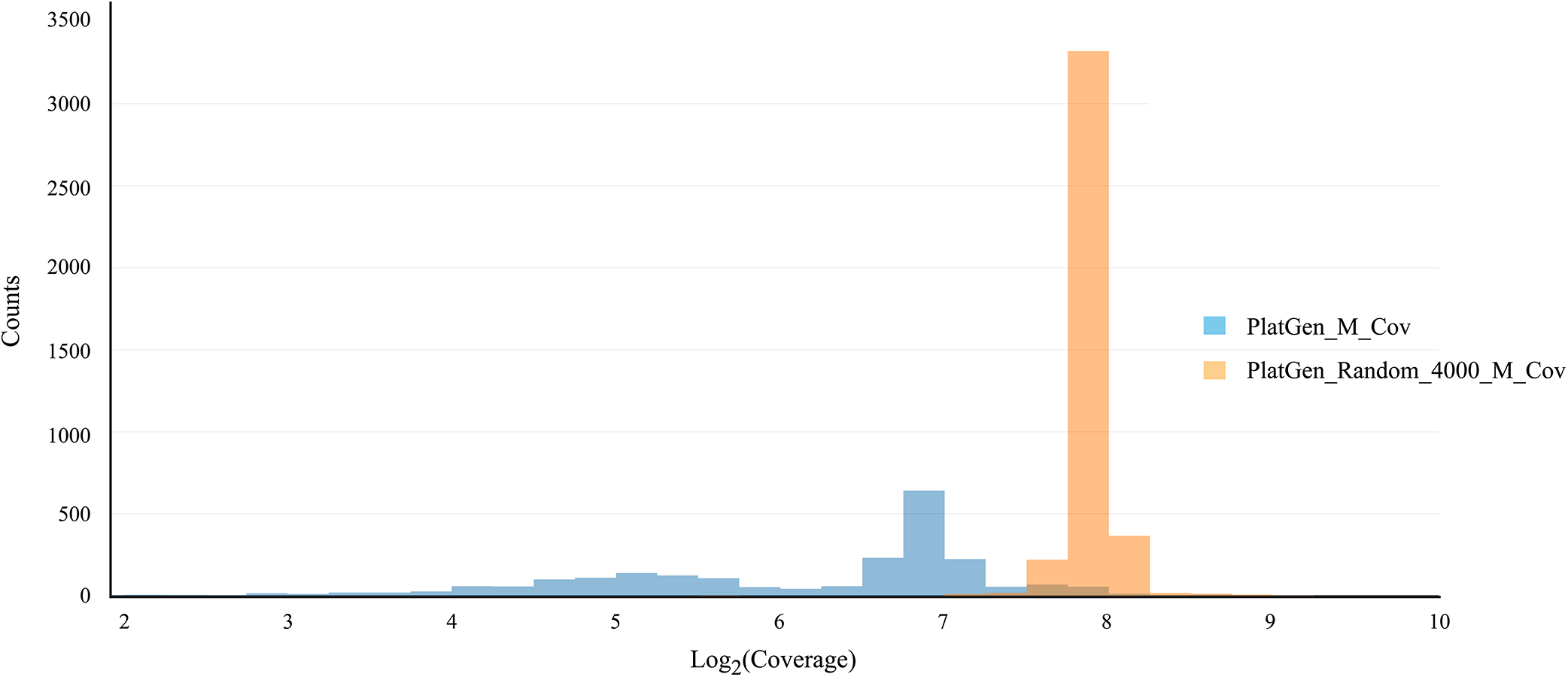
Depth of coverage distribution for Personalis deletion calls (PlatGen_M_Cov) and random regions (PlatGen_Random_4000_M_Cov). See original data at https://plot.ly/337/~parikhhm/.

**Figure 3:**
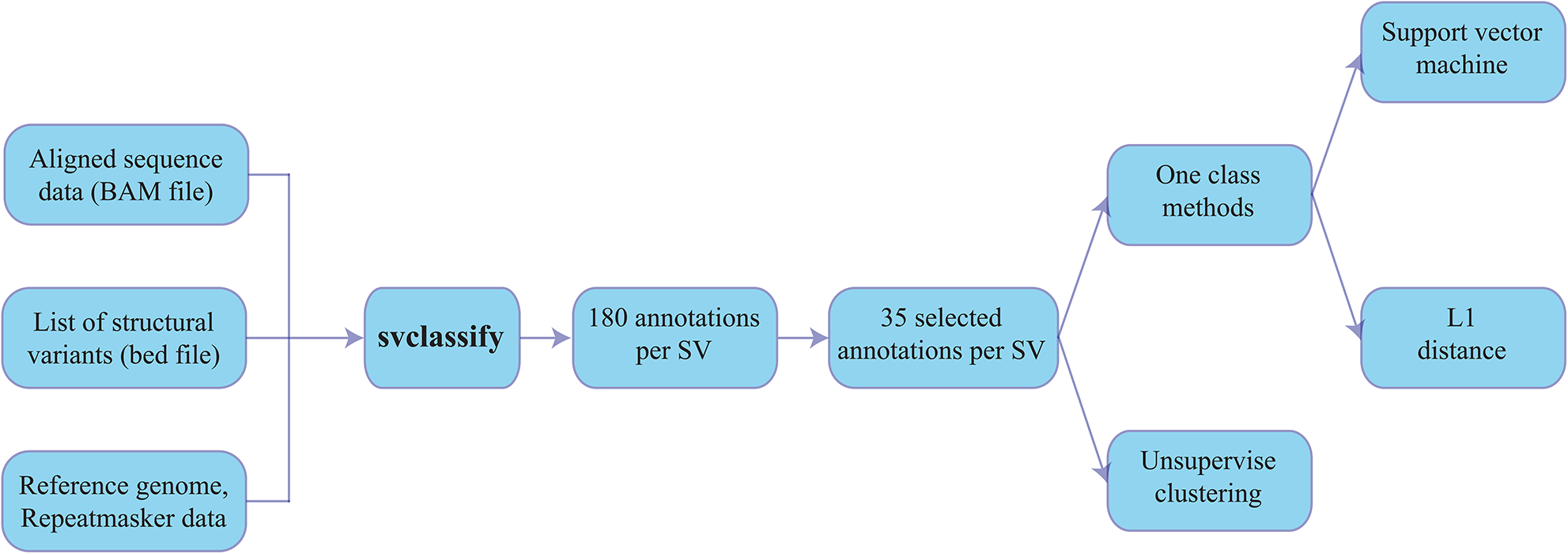
Flowchart of analytical approach to classify candidate SVs into likely true or false positives. The subset of 35 annotations was chosen for Illumina paired-end data (fewer for PacBio and moleculo data) to reduce the number of annotations used in the model to those that we expected to be most important for clustering calls into different categories. The one-class model uses only the 4000 random sites for training, and it assumes that sites with annotations unlike most of these random sites are more likely to be SVs.

For annotations for one-class classification, we included the coverage inside the SV region (M_Cov, LM_Cov, and RM_Cov) because coverage should be directly proportional to copy number. We included soft-clipping in the flanking regions because we usually expect only part of reads to map when they cross the SV breakpoints. We included insert size in the flanking regions because we expect insert size to be larger than normal for deletions and smaller than normal for insertions. We included the proportion discordantly mapped reads because this can indicate either a large insert size or a mobile element insertion. We included the proportion of reads for which the other end is not mapped because this could indicate a novel insertion. For PacBio, we included the difference between inserted and deleted bases because the reads were often sufficiently long to be mapped across a deleted or inserted region and abnormally high insertion or deletion rate can indicate an insertion or a deletion event.

For clustering, we included additional annotations that may not distinguish true SVs from non-SVs but could be helpful for clustering different types of true SVs. For example, low mapping quality might indicate a homozygous deletion, a mobile element deletion, or a biased non-SV region. Also, SV size could be useful to cluster different types of SVs. Abnormally low or high coverage in the flanking regions could indicate imprecise breakpoints, regions with bias, or a much larger SV encompassing the region. We also only used for clustering the reference-specific or sample-specific annotations like RepeatMasker regions, GC content, and number of heterozygous and homozygous SNPs. We did not include these in the one-class model because they are not technology-specific, so they could not simply be used as part of our heuristics requiring evidence from multiple technologies.

### Hierarchical cluster analysis separates random regions from multiple SV types

To understand the types of SV calls in the validated/assembled deletion sets and how they segregate from random genomic regions, we first performed unsupervised machine learning using hierarchical clustering with a manually selected subset of 11 to 35 annotations from svclassify, depending on the technology (Supplementary table 1 to 4). This subset of annotations was chosen to reduce the number of annotations used in the model to those that we expected to be most important for clustering calls into different categories. We decided to focus our analyses on eight major clusters, which are visualized as a tree (dendrogram) in Figure 4A and with multidimensional scaling in Figures 4B and 4C. Five of the clusters (1, 2, 3, 6, 7) were predominantly (98.5 %) SVs, two clusters (4 and 5) were predominantly (98.9 %) non-SVs, and one cluster (8) was 40 % SVs and 60 % non-SVs. The label (SV or non-SV) associated with each site was not provided to the clustering method, and yet the clusters showed a good separation of SVs from non-SVs based entirely on the annotation values. To ensure the 4000 random regions sufficiently represented non-SVs, we also generate random regions matching the size distribution of the Personalis deletions, as well as random SINEs, LINEs, and LTRs. It is promising that even the randomly selected SINEs, LINEs, and LTRs generally segregate with the random genomic regions even though they are from regions of the genome that are difficult to map.

**Figure 4:**
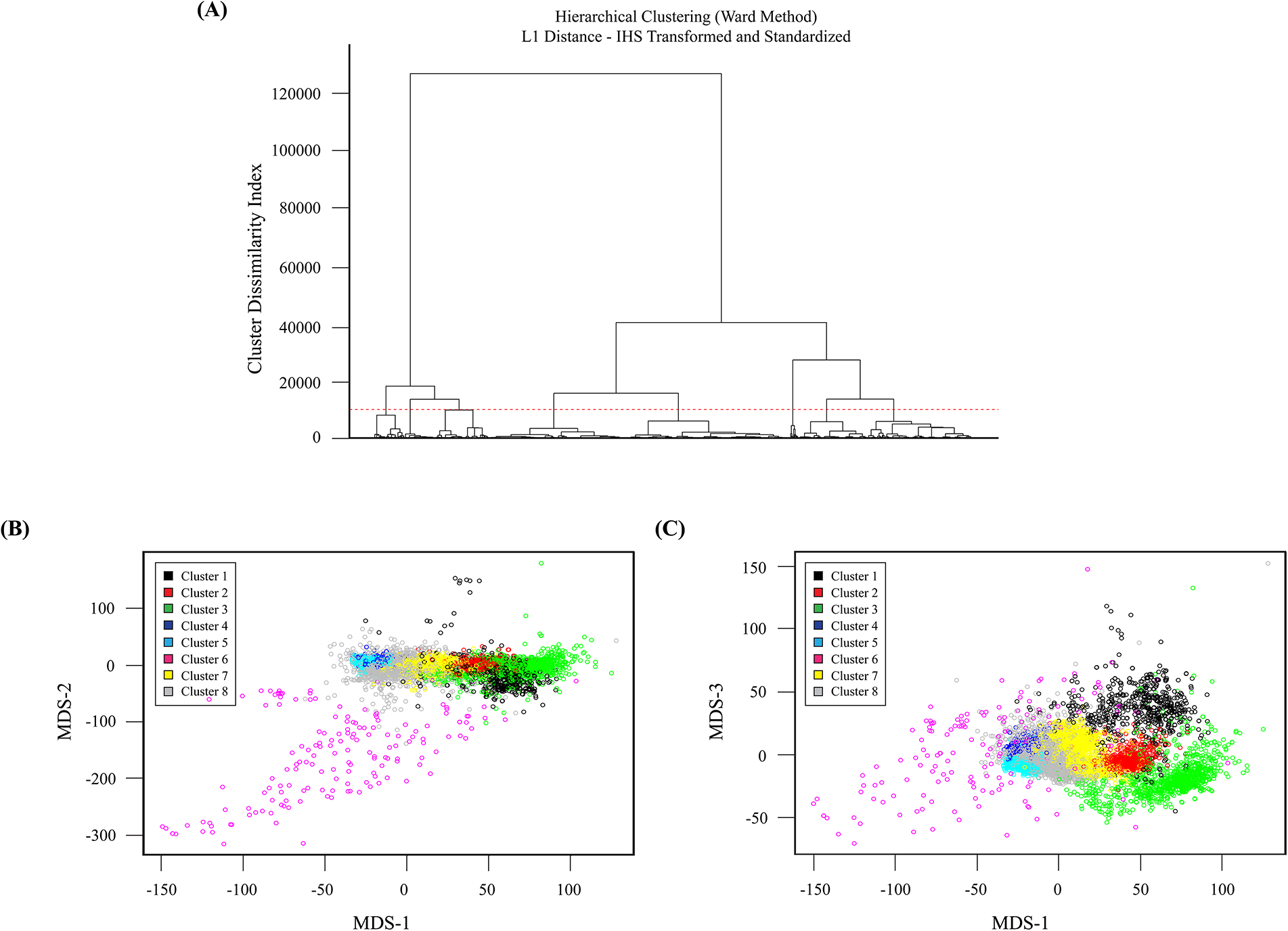
Hierarchical clustering results using L_1_ distance and Ward’s method shown as (A) a dendrogram and (B-C) in multi-dimensional scaling plots. (A) The horizontal dotted red line shows the cut-off at a cluster dissimilarity index of about 10000, which results in 8 clusters. The clusters are number 1 to 8 from left to right, with 4 and 5 containing primarily non-SVs, 8 containing a mixture of SVs and non-SVs, and 1, 2, 3, 6, and 7 containing different types of deletions (see Table 4). (B-C) Multidimensional scaling plots for visualizing the 8 clusters. We use a 3 dimensional representation of the data space which associates 3 MDS coordinates to each site, one for each dimension. (B) Plot of MDS-2 against MDS-1, which clearly separates Cluster 6 (mainly SVs with inaccurate breakpoints). (C) Plot of MDS-3 against MDS-1, in which the different types of SVs are generally well-separated from each other and from non-SVs.

We further compared the annotations of these 8 clusters to understand whether they represent different categories of SVs and random regions. Clusters 4 and 5 contain close to 99 % non-SVs, but Cluster 4 generally contains larger sites than Cluster 5. Cluster 8 is a mix of 60 % non-SVs and 40 % SVs, and sites in Cluster 8 generally have a coverage between the normal coverage and half the normal coverage, and more sites have lower mapping quality, repetitive sequence, and high or low GC content. Further subdivisions of Cluster 8 might divide the true SVs from non-SVs.

98.5 % of sites in Clusters 1, 2, 3, 6, and 7 are from the Personalis and 1000 Genomes Gold sets, but the clusters contain different types of SVs. Clusters 1, 2, 3, and 6 generally contain reads with lower mapping quality inside the SV, though the low mapping quality could arise from a variety of sources (e.g., repetitive regions that are falsely called SVs, true heterozygous or homozygous deletions of repetitive elements like Alu elements, or true homozygous deletions that contain some incorrectly mapped reads inside the deletion). Clusters 2 and 3 appear to be true deletions of Alu elements, since sites in these clusters are ~300 bps, are annotated as SINEs, LINEs, or LTRs by RepeatMasker, have high GC content, and have low mapping quality. Cluster 2 sites are primarily heterozygous Alu deletions since they have about half the typical coverage, and Cluster 3 sites are primarily homozygous Alu deletions and a small fraction of other homozygous deletions because they contain less than half the typical coverage. All 655 sites in Cluster 1 are from Personalis and 1000 Genomes, and appear to be mostly larger homozygous deletions (half are larger than 2000 bps), and they have lower than half the normal coverage, low mapping quality, and more discordantly mapped reads. 86 % of sites in Cluster 6 are from 1000 Genomes and appear likely to represent mostly true homozygous deletions with imprecise breakpoints that are too narrow, since the left and right flanking regions, in addition to the region inside the putative SV, have low coverage less than half the typical coverage. 97.4 % of sites in Cluster 7 are from Personalis and 1000 Genomes, and they appear to be predominantly heterozygous deletions in relatively easier parts of the genome with high mapping quality. These results are summarized in Table 3.

**Table 3:** Analysis of 8 clusters from hierarchical cluster analysis, including the numbers of sites from each call set and a description of the predominant types of sites in each cluster.

| Cluster | 4000<br>Random | Personalis<br>Random | Random<br>LINEs | Random<br>LTRs | Rand<br>om<br>SINEs | Personalis<br>deletions | 1000<br>Genomes<br>deletions | Total | Proporti<br>on of<br>deletions | Description |
| --- | --- | --- | --- | --- | --- | --- | --- | --- | --- | --- |
| 1 | 0 | 0 | 0 | 0 | 0 | 371 | 284 | 655 | 1.000 | Mostly large,<br>true<br>homozygous<br>deletions |
| 2 | 0 | 0 | 0 | 0 | 2 | 432 | 237 | 671 | 0.997 | Heterozygous<br>Alu deletions |
| 3 | 1 | 1 | 1 | 0 | 0 | 705 | 402 | 1110 | 0.997 | Homozygous<br>Alu deletions |
| 4 | 2397 | 455 | 38 | 28 | 16 | 9 | 28 | 2971 | 0.012 | Large, likely<br>non-SVs.<br>Generally in<br>easy-to-<br>sequence<br>regions |
| 5 | 1073 | 1351 | 352 | 378 | 279 | 1 | 33 | 3467 | 0.010 | Smaller,<br>likely non- |
|  |  |  |  |  |  |  |  |  |  | SVs.<br>Generally in<br>easy-to-<br>sequence<br>regions |
| 6 | 17 | 2 | 1 | 0 | 0 | 3 | 138 | 161 | 0.876 | Likely true<br>large<br>homozygous<br>deletions with<br>inaccurate<br>breakpoints<br>so that the<br>true deletion<br>is larger than<br>the called<br>region |
| 7 | 14 | 16 | 2 | 2 | 4 | 624 | 811 | 1473 | 0.974 | Mostly true<br>heterozygous<br>deletions in<br>easier-to-<br>sequence<br>regions |
| 8 | 498 | 481 | 103 | 90 | 195 | 161 | 752 | 2280 | 0.400 | Mix of non-SVs and SVs in more difficult regions with coverage between the normal coverage and half the normal coverage |
| **Total** | 4000 | 2306 | 497 | 498 | 496 | 2306 | 2685 | 1278<br>8 | 0.390 |  |

More sophisticated versions of our clustering approach are available. Parametric approaches include Gaussian mixture modeling, but there are also nonparametric mixture modeling approaches available. However, exploratory analyses showed that at best only a marginal improvement is realized using such more advanced methods for our datasets.

### One-class classification of candidate SVs using L_1_ distance integrates information from multiple technologies

We next developed a one-class classification model to classify candidate sites as high-confidence SVs or uncertain. This one-class model uses only the 4000 random sites for training, and it assumes that sites with annotations unlike most of these random sites are more likely to be SVs. As shown in Supplementary table 5, we only used a subset of the annotations from the unsupervised hierarchical clustering because atypical values for some annotations (e.g., mapping quality or SV size) do not necessarily indicate that an SV exists in this location (see the section above about annotations for why these annotations were selected). The number of annotations used ranged from 7 for PacBio to 30 for Illumina paired-end because certain annotations like insert size do not apply to all technologies (Supplementary table 1 to 4).

Results from the L_1_ distance one-class classification are summarized using ROC curves. Five different ROC curves are shown in Figure 5A-5B, one from each classifier using one of the four data sets and one classifier based on all datasets combined. The classifier based on all datasets combined performs the best with PlatGen (200x Illumina) alone being a close second. ROC curves for the ensemble classifiers, based on the four L_1_ classifiers using each of the four data sets separately, are shown in Figure 5C-5D. Four different ensemble classifiers are considered based on four different ways of combining the results from the individual classifiers. A typical ensemble classifier will classify a site as SV if k or more of the individual classifiers make an SV call. Here k can be 1, 2, 3, or 4. The results show that using k=3 provides the best ensemble classifier with k=2 being a close second. Performance is similar for the k=3 classifier and all datasets combined, and we use k=3 for our final results because we expect requiring evidence from 3 datasets will be more robust. For k=3, we calculated the proportion ρ of random sites that are closer to the center than each candidate site. We stratified candidate sites into those with ρ < 0.68, 0.68 < ρ < 0.9, 0.9 < ρ < 0.99, or ρ > 0.99, as shown in Table 4.

**Table 4:** Number of sites from each candidate call set that have k=3 L_1_ Classification scores in each range, where the score is the proportion ρ of random sites that are closer to the center than each candidate site. These numbers are after filtering sites for which the flanking regions have low mapping quality or high coverage.

|  | **Filtered** | **<0.68** | **0.68-0.9** | **0.9-0.97** | **0.97-0.99** | **0.99-0.997** | **0.997-0.999** | **>0.999** |
| --- | --- | --- | --- | --- | --- | --- | --- | --- |
| Random<br>Personalis | 229 | 3025 | 501 | 177 | 65 | 3 | 0 | 0 |
| Personalis | 106 | 8 | 10 | 44 | 414 | 409 | 1302 | 13 |
| Gold |  |  |  |  |  |  |  |  |
| Personalis<br>Validated | 3 | 0 | 0 | 0 | 10 | 7 | 19 | 0 |
| Personalis<br>Non-validated | 0 | 1 | 0 | 0 | 3 | 0 | 1 | 0 |
| 1000<br>Genomes | 382 | 56 | 103 | 257 | 714 | 388 | 780 | 5 |
| Spiral Gen<br>Insertions | 1 | 0 | 0 | 0 | 12 | 16 | 41 | 0 |
| Deduplicated<br>Deletions | 195 | 45 | 61 | 145 | 675 | 513 | 1434 | 14 |

**Figure 5:**
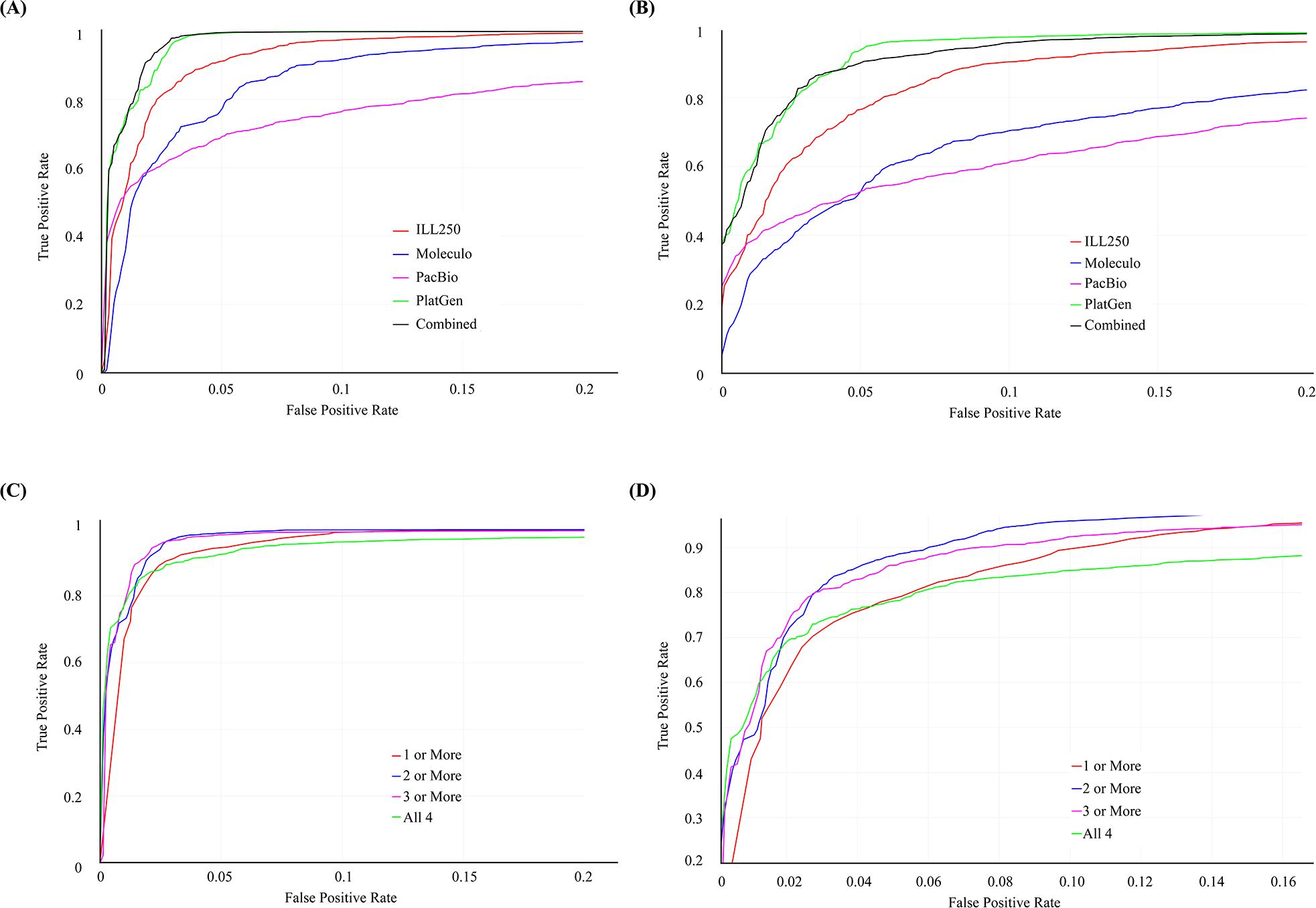
ROC curves for One-class classification using the L_1_ Distance, treating the 4000 Random regions as negatives and the Personalis or 1000 Genomes calls as positives. (A) ROC curves for one-class models for each dataset separately and for all combined for the Personalis validated deletion calls. (B) ROC curves for one-class models for each dataset separately and for all combined for the 1000 Genomes validated deletion calls. (C) ROC curves for one-class model requiring 1 or more, 2 or more, 3 or more, or all 4 technologies to have high classification scores for the Personalis validated deletion calls. (D) ROC curves for one-class model requiring 1 or more, 2 or more, 3 or more, or all 4 technologies to have high classification scores for the 1000 Genomes validated deletion calls. The 3 or more classification method is used to produce the final high-confidence SVs in this work. The horizontal axis shows the false positive rate (from the random set of regions matching the size distribution of the Personalis deletions) and the vertical axis shows the corresponding true positive rate (assuming all the validated/assembled calls are true). See original data at https://plot.ly/345/~parikhhm/, https://plot.ly/353/~parikhhm/, https://plot.ly/361/~parikhhm/, and https://plot.ly/369/~parikhhm/.

To confirm our choice of 4000 random regions for training, we compared the scores of the 4000 random regions to the 2306 different random regions with a size profile matching the Personalis SVs. We found that the distributions of scores were very similar, with 23 of 2306 Personalis random regions having ρ > 0.99, close to the 1 % expected. Similarly, 11 of 1491 random SINE, LINE, and LTR regions had ρ > 0.99, close to the 1 % expected. To assess the performance of the classifier on random regions, we also found the distribution of ρ scores for the random Personalis regions when assessed against the 4000 random regions, and the distribution of scores was flat as expected (see Supplementary figure 1).

### One-class classification of candidate SVs using SVM gives similar results to L_1_ classifier

To compare to an alternative distance measure and method for one-class classification, we also developed a one-class SVM model. We found that results were generally similar between the L_1_ one-class results and the SVM one-class results in terms of ROC curves (Supplementary figures 2, 3, 4, and 5). Supplementary table 6 gives the concordance/discordance matrix for predictions from the L_1_ and SVM one-class classifications for selected values of ρ. Agreement between the two methods is 84 % with ρ > 0.99, 98 % with ρ > 0.95 and 99 % with ρ > 0.9, on Personalis validated/assembled set. The high agreement between SVM and L_1_ at ρ > 0.95 suggests that our one class classification method is robust to the type of model. We further examined the 7 sites consistently identified with only SVM and 1 site consistently identified with only L_1_ that had low ρ (0.6 > ρ > 0.5) with one method and ρ > 0.9 with the other method. We found that these were from difficult regions of the genome, such as telomeres, high coverage regions, and low mapping quality regions, so they are filtered from our final high-confidence calls. However, similar comparisons of predictions on 1000 Genome set with L_1_ and SVM ensemble classifiers suggest that the L_1_ classifier has better efficiency in predictions on 1000 Genome set and better agreement on different technologies. Therefore we use the simpler L_1_ method.

### Manual inspection of one-class results verifies accuracy of our classifier

We randomly selected a subset of sites from each call set in each selected ρ value range from Table 4 for manual inspection. In general, Personalis and 1000 Genomes sites with high ρ values were very likely accurate and mostly homozygous, while sites with lower ρ appeared to be questionable, small, and/or heterozygous. Most of the Spiral Genetics insertions had very high ρ, indicating a true SV is likely in the region.

For Personalis, we inspected 20 randomly selected sites with ρ > 0.99, and all appeared to be accurate (Supplementary table 7). Only 5 of the 20 sites appeared likely to be heterozygous, since homozygous deletions generally are more different from random regions than heterozygous deletions. 4 out of 5 heterozygous sites had 0.99 < ρ < 0.999, whereas all 15 homozygous deletions had ρ > 0.999 except for one small 52-bp deletion. 13 of the homozygous deletions had ρ > 0.9999. Also, all 10 of the randomly selected Personalis sites with 0.9 < ρ < 0.99 were likely to be true heterozygous deletions, and none were homozygous (Supplementary table 8). There were only 8 sites with ρ < 0.9 in the Personalis set (Supplementary table 9), and these were a mixture of likely true but very small deletions and other potential deletions that were difficult to determine whether they were true or artifacts since they were only supported by a small number of reads. Therefore, we do not include these calls with ρ < 0.9 in our final high-confidence set.

For 1000 Genomes, we similarly inspected 20 randomly selected sites with ρ > 0.99, and all appeared to be accurate except for one in a low complexity region, which had few supporting reads in svviz. Only 4 (20 %) of the sites with ρ > 0.99 had ρ > 0.9999, in contrast to 65 % of the Personalis calls. 3 of the 4 sites with ρ > 0.9999 were likely to be homozygous deletions. One likely true heterozygous deletion had ρ > 0.999, and the remaining 15 sites with 0.99 < ρ < 0.999 appeared likely to be true heterozygous deletions except for one in a low complexity region (Supplementary table 10). Also, 7 of the 9 randomly selected 1000 Genomes sites with 0.9 < ρ < 0.99 were likely to be true heterozygous deletions, and none were homozygous (Supplementary table 11). The other 2 sites contained 17 % and 58 % low complexity sequence and 68 % and 66 % GC content, and they appeared likely to be erroneous calls since no reads aligned to the alternate allele for any technology using svviz (except for a single moleculo read for one of the sites). 7 of the 8 randomly selected 1000 Genomes sites with 0.7 < ρ < 0.9 were smaller than 100 bps, 6 were likely to be true heterozygous deletions, and none were homozygous (Supplementary table 12). 5 of the 7 randomly selected 1000 Genomes sites with ρ < 0.7 were smaller than 110 bps and were possibly true heterozygous deletions, and none were homozygous (Supplementary table 13). In general, the 1000 Genomes calls have lower ρ scores than the Personalis calls because the Personalis calls contain a higher fraction of homozygous deletions, fewer very small deletions, and are all breakpoint-resolved.

All of the complex insertions from Spiral Genetics had ρ > 0.97, indicating that they are likely to be true SVs. Upon manual inspection of the svviz results (Supplementary table 14), 29 had evidence in all 4 technologies for a homozygous insertion, 29 had evidence in all 4 technologies for a heterozygous insertion, and 8 were inconsistent in terms of zygosity across the 4 technologies. The reason for the discordance between technologies for the 8 discordant sites is not always clear, but it appears that some are likely to be real SVs with different breakpoints. For example, an insertion is called at 1:3,418,563 with a length of 352 bp, but appeared likely to be much larger.

Most candidate sites with ρ > 0.9 appear to be true, but a few of the manually inspected sites appeared to be inaccurate or to have incorrect breakpoints. Therefore, we further refined our final call set by using svviz to map reads to the reference or predicted alternate alleles, and we included only sites with at least 3 reads supporting the alternate allele in at least 3 of the 4 datasets. This filtered 13 % percent of the calls, leaving 2676 deletions and 68 insertions for which we have high confidence. These calls are publicly available at ftp://ftp-trace.ncbi.nih.gov/giab/ftp/technical/svclassify_Manuscript, and we will continue to update these with additional call sets as we further develop our methods.

### PCR validates high-confidence SVs

To obtain estimates of accuracy of the Personalis deletion calls, we performed experimental validation for some of the calls. Only 44 of 2350 calls met the criteria for designing primers, 3 primer pairs failed, and in one case we were unable to make a call. We were able to validate 38 of Personalis’ deletions with exact breakpoints (including 3 within 1 bps) out of the 40 deletions that we could test. A 39th case was off by 44 bps on one side and the last case was a false positive call. All homozygous calls (6) were confirmed by the validation. Only 10 out of 21 heterozygous calls had the correct zygosity call. Of the heterozygous calls with incorrect zygosity, 7 were actually homozygous, 1 could not be determined by the validation and 1 was not a deletion. The remaining cases did not have a zygosity call, of which 9 were homozygous and 7 were heterozygous. All of the validated calls had ρ > 0.97. 4 of the 5 sites that could not be PCR-validated also had ρ > 0.97. The fifth site (chr9:8860664-8875135), which was the only likely false positive, had ρ = 0.01 and is bounded by approximately 350 bp regions that only differ by a 9 bp indel.

### Trio analysis with MetaSV confirms most high-confidence SVs

Given the rate of *de novo* mutation is low [11], most SVs of an individual are expected to be Mendelian consistent. We went on to assess the quality of our high-confidence SVs by validating it with trio analysis, involving the child (NA12878), her father (NA12891) and her mother (NA12892). Sequences were retrieved from the Illumina Platinum Genomes repository with an average sequencing depth of approximately 50x. MetaSV [5], a recently published SV caller integrating multiple orthogonal SV detection algorithms, was used to generate three call sets for the trio individuals. For highest quality, only deletion calls >=100 bps were considered. Calls detected by two or more algorithms in MetaSV with different detection methods were deemed as PASS and regarded as high-quality [5, 12]. There were 13639, 34565, and 32482 deletion calls for NA12878, NA12891, and NA12892 respectively, of which 2671, 2714, and 2640 were PASS calls.

To validate the 2676 high-confidence deletion SVs from svclassify, we selected the 2348 deletions >= 100 bps and matched them with the Bina call sets. We used a 50% reciprocal overlapping strategy for the matching. A call from svclassify was deemed as “validated” if one of the following criteria was met: Level (1) detected in any of the two parents from the Bina call sets; Level (2) detected in the child as a PASS from the Bina call sets; and Level (3) reported and validated in previous literatures on the child [13]. As shown on Supplementary figure 6, 98% of the high-confidence deletions were confirmed by the Level 1 validation, i.e., also detected in the parents. There were an additional 0.2% confirmed by the Level 2 validation, i.e., detected by at least two orthogonal algorithms. Level 3 validation confirmed an extra 1.5%, resulting in a total validation rate of 99.7%. The extremely high validation rate indicates high quality of the high-confidence SV call set from svclassify.

### Performance of svclassify with 30x coverage dataset

To understand the effect of downsampling on the classification accuracy, we performed downsampling of the Platinum Genomes BAM file to 30x mean coverage. As shown on Supplementary figure 7, even with 30x coverage, at a 5% false positive rate, > 97 % of the Personalis calls were classified as true positives, but this is lower than the >99 % of calls classified as true positives at a 5% false positive rate with 200x coverage.

## Conclusion

High-confidence SV and non-SV calls are needed for benchmarking SV callers. To establish high-confidence, methods are needed to combine multiple types of information from multiple sequencing technologies to form robust high-confidence SV and non-SV calls. Therefore, in this work we developed methods to classify SVs as high-confidence based on annotations calculated for multiple datasets. Our classification method gives the highest scores to SVs that are insertions or large homozygous deletions, and have accurate breakpoints. Deletions smaller than 100-bps often have low scores with our method, so other methods like svviz are likely to give better results for very small SVs. Homozygous deletions generally receive the highest scores because they have annotations most unlike random regions of the genome. Breakpoint-resolved deletions generally receive higher scores because reads near the breakpoint have distinct characteristics such as clipping and insert size that our method uses to classify SVs. We produce a set of 2676 high-confidence deletions and 68 high-confidence insertions with evidence from 3 or more sequencing data sets. These sets of SVs are likely biased towards easier regions of the genome and do not contain more difficult types of SVs. However, they can be used as an initial benchmark for sensitivity for deletions and insertions in easier regions of the genome.

In this work, we use data from multiple whole genome sequencing technologies to develop high-confidence SVs for benchmarking, and it is important to understand strengths and weaknesses of this approach. Alternative approaches might include targeted experimental validation and simulation of SV events. For Reference Materials that are characterized by multiple high-coverage whole genome sequencing technologies, we expect our approach to work well, though it may still be useful to confirm some SVs with targeted methods. It is sometimes difficult or impossible to target SVs in difficult regions of the genome, so it is important to understand that this type of confirmation may not help assess the accuracy of the most difficult SVs. Similarly, simulated SVs in random regions of the genome may not represent the true locations of SVs, which can often occur in challenging regions of the genome and may have sequencing errors that are not simulated. While SVs with more accurate breakpoints are more likely to have high scores with our method, one or both breakpoints may be inaccurate for some high-confidence SVs, and even when accurate it is often possible to represent SVs in multiple ways in repetitive regions. It is also critical to understand that the high-confidence SVs in this manuscript likely represent a subset of SVs biased towards those that are easier to detect and are generally moderate sized insertions and deletions. In addition, although multiple SV callers and analysis groups generated the candidate calls, they may be biased towards existing callers and short-read data. Therefore, sensitivity measured with respect to our high-confidence SVs or most other high-confidence SVs is likely an overestimate of the sensitivity of any method for all SVs in the genome. Further work is needed to develop more comprehensive high-confidence SV call sets.

The motivation for building the one-class model using random regions rather than true SVs is that we do not have unbiased sets of true SVs that we could use for training. Unfortunately, current “true SV” call sets tend to be biased towards types of SVs that are easier to call and in easier regions of the genome. For example, a model based on moderate-sized deletions is unlikely to be useful for insertions, duplications, large or small deletions, or SVs in more difficult parts of the genome. A one-class model based on true SVs also may result in overfitting, which is mitigated by using random regions because the model is generated independent of the true SVs used in validation. A limitation of using random regions is that they may not perfectly represent false positive SVs that are generated by callers. Further work will be needed to understand the profiles of false positive SVs and how they compare to random regions, since we did not have sufficient false positives to test this in this work.

The primary goal of this work is to develop methods to form high-confidence SV calls from candidates generated from multiple technologies and calling methods. However, future work might also examine using these methods to classify candidate SV calls from multiple callers using a single technology. The utility of svclassify in this case would be to generate and use consistent annotations for all candidate SVs. While multiple technologies are likely to result in more robust classifications, even a single technology often separates most of our high-confidence SVs from random regions.

Our unsupervised clustering methods also show promise for classifying candidate SVs into different types and potentially classifying more difficult types of SVs. Seven of the eight clusters obtained from an unsupervised hierarchical cluster analysis using L_1_ distances were relatively pure clusters consisting of either mostly SVs or mostly non-SVs. The overall successful separation of the SVs from the non-SVs by the unsupervised analysis suggests that the annotations for SVs and non-SVs occupy more or less disjoint regions in the data space. Since each cluster contains a different type of SV or non-SV, future work might include further investigation of these clusters and sub-clusters to understand their meaning. In addition, we plan to apply these clustering methods to additional types of SVs and develop more sophisticated classification methods that would place new candidate SVs in one of these categories of different types of true or false positive SVs.

We plan for the methods developed in this work to form a basis for developing high-confidence SV and non-SV calls for the well-characterized NIST RMs being developed by the GIAB. In this work, we apply these methods to produce a set of high-confidence deletions and insertions with evidence from multiple sequencing datasets, and we plan to continue to develop these methods to be applied to more difficult types of SVs in more difficult regions of the genome. We also plan to incorporate calls from methods merging multiple callers, such as MetaSV [5]. For example, there were 456 high-quality MetaSV PASS calls for NA12878 which were also Mendelian consistent. Finally, we hope to incorporate statistics from other tools, such as Parliament [9] and svviz [10], in our machine learning models.

## Materials and Methods

### Data sets

Four whole-genome sequencing data sets (Table 1) were used to develop methods to classify candidate SVs into true positives and false positives for Coriell DNA sample NA12878. Two data sets were generated using short-read sequencing technologies, and two other data sets were generated using long-read sequencing technologies. For the Platinum Genomes 2x100bps HiSeq data, raw reads were mapped to the National Center for Biotechnology Information (NCBI) build 37 using the Burrows-Wheeler Aligner (BWA) “bwa mem” v.0.7.5a with default parameters [14]. For Illumina HiSeq (read length = 250 bps), PacBio, and Moleculo whole-genome sequencing data sets, aligned bam files were publicly available and were used directly in this study.

### SV validated/assembled sets

Three validated/assembled SV sets (Table 2) totaling 5035 deletions and 70 insertions were derived from Coriell DNA sample NA12878.

(A) Personalis deletions calls were derived based on pedigree analysis, which included 16 members of the family.

To be included in the validated/assembled set, the following conditions had to be met:

1. Deletion must have been detected in at least one NA12878 sample.
2. Deletion must have been detected in at least 2 other samples in the pedigree with exact breakpoint matches.

The Personalis gold data set was further refined by experimental validations. Primers were designed based on following criteria:

1. Each primer maps no more than 3 times in genome.
2. Require unique polymerase chain reaction (PCR) product in genome.
3. 400 – 800 bps product size.
4. Pad 100 bps around each deletion junction.

For small deletions (< 200 bps) a single primer pair was designed that straddled the deletion. For large deletions (> 1500 bps) two primer pairs were designed around each reference breakpoint junction. Site specific PCR amplification and high depth MiSeq shotgun sequencing followed by manual inspection of the alignments was used to validate all the deletions. Sanger sequencing was used when we were not able confirm the deletion with MiSeq. For 3 deletions (2:104186941-104187136, 7:13022102-13028550, and 14:80106289-80115049) this was done because we did not see any junction reads.

(B) The 1000 Genomes Project validated/assembled contains the set of validation deletion calls found in the genome of NA12878 by the 1000 Genomes Project pilot phase [13, 15]. These deletion calls were validated by assembly or by other independent technologies such as array comparative genomic hybridization, sequence capture array, superarray, or PCR.

(C) Spiral Genetics’ Anchored Assembly was performed whole read overlap assembly on corrected, unmapped reads to detect structural variants using Illumina 2x100bps HiSeq whole-genome sequencing data set. Sequencing errors were corrected by counting k-mers. Low count k-mers were discarded as erroneous. The set of high scoring, or true k-mers was used to construct a de Bruijn graph representing an error-free reconstruction of the true read sequences. Each read was corrected by finding the globally optimum base substitution(s) so that it aligned to the graph with no mismatches and differed by the smallest base quality score from the original read. Of these corrected reads, those that did not match the reference exactly were assembled into a discontiguous read overlap graph to capture sequence variation from the reference. Variants were mapped to human reference coordinates (NCBI build 37) by walking the read overlap graph in both directions until an “anchor” read, where a continuous 65 bps matches the reference, denoted the beginning and end of each variant. Where a variant had more than one anchor, pairing information was used to determine the correct location of the anchor. We used 70 calls from the “Insertions” output, all of which were complex insertions (i.e., a set of reference bases was replaced by a larger number of bases).

### Deduplicated deletions

Any overlapping deletions within the validated/assembled SV sets were discarded because they may represent compound heterozygous SVs or imprecise or inaccurate SVs. This deduplication resulted in 2336 unique Personalis deletion calls and 1825 unique 1000 Genomes deletion calls (Table 2). Bedtools’ intersect function was used to screen overlap between these two datasets (Supplementary table 15). Merged deduplicated deletion calls were generated by keeping all the 2336 unique Personalis deletion calls and merging with 746 non-overlapping 1000 Genomes deletion calls with minimum overlap required to be 1 bp, which resulted 3082 deduplicated deletion calls.

### Random region non-SV call sets

In addition, five sets of likely non-SVs were generated: 2 random and 3 from repetitive regions of the genome (Table 2) as follows:

1. 4000 random regions were generated with a uniform size distribution on a log scale from 50 bps to 997527 bps. Start sites were chosen randomly using the Generate Random Genomic Coordinates script in R (http://www.niravmalani.org/generate-random-genomic-coordinates/).
2. 2306 random regions were generated with a size distribution matching the calls from the pedigree-based Personalis deletions call set. Start sites were chosen randomly using the Generate Random Genomic Coordinates script in R (http://www.niravmalani.org/generate-random-genomic-coordinates/).
3. 497 long interspersed nuclear elements (LINEs) were randomly selected from a list of LINEs from the University of California, Santa Cruz (UCSC) Genome Browser’s RepeatMasker Track.
4. 498 long terminal repeat elements (LTRs) were randomly selected from a list of LTRs from the UCSC Genome Browser’s RepeatMasker Track.
5. 496 short interspersed nuclear elements (SINEs) were randomly selected from a list of SINEs from the UCSC Genome Browser’s RepeatMasker Track.

### svclassify

The svclassify tool was developed to quantify annotations of aligned reads inside and around each SV (Figure 1). It was written using the Perl programming language employing SAMtools (version 0.1.19-44428cd) [16] and BEDTools (version 2.17.0) [17] to calculate parameters such as coverage, paired-end distance, soft clipped reads, mapping quality, numbers of discordant paired-ends reads, numbers of heterozygous and homozygous SNP genotype calls, percentage of the GC-content, percentage of the repeats and low complexity DNA sequence bases, and mapping quality. svclassify requires the following inputs: a BAM file of aligned reads, a list of SVs, homozygous and heterozygous SNP genotype calls, a list of repeats from the UCSC Genome Browser’s RepeatMasker Track and a reference genome. BAM files can come from any aligner. The user can specify the size for the flanking regions. svclassify also includes partially mapped reads to the L, LM, M, RM, or R regions for calculations. The insert size is calculated as the end-to-end distance between the reads (length of both reads + distance separating the reads). Because PacBio reads have high insertion and deletion error rates, Del (the mean of deleted bases of the reads) and Ins (the mean of inserted bases of the reads) were normalized by subtracting the mean Del (0.0428) and Ins (0.0948) per read length of 4000 random regions. For exploratory analyses, svclassify generates 85 to 180 annotations for each SV from each dataset, depending on sequencing technology (Supplementary table 5 and 16). For our unsupervised and one class analyses, we used only subsets of these annotations that we expected to give the best results. These subsetted annotations are given in the csv files (Supplementary table 1 to 4).

### Data Analysis

The results from svclassify were subjected to two types of analyses – (1) Unsupervised Learning based on a hierarchical cluster analysis using the L_1_ distance (also called *Manhattan distance*), and (2) One Class Classification using the L_1_ distance or support vector machines (SVM) using a carefully selected set of 4000 non-SVs.

### Unsupervised Learning

Data values for each variable (characteristic) used in the analysis were first transformed using an inverse hyperbolic sine transformation [18]. This transformation uses the following function.

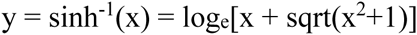

This function is often used as an alternative to the logarithmic transformation. It has the advantage that zero or negative values of x do not cause problems. Generally speaking it is quite similar to a standard logarithmic transformation except near and below zero. Next, all variables were standardized by subtracting the mean and dividing by the standard deviation. All further work was done using these transformed data.

A hierarchical cluster analysis was performed with all 7797 random sites, 5035 deletion sites, and 70 insertion sites (see Table 2), using L_1_ distance as the distance function rather than Euclidean distance [19] since Manhattan distance is less influenced by outliers within the non-SV class. The Ward method was used for clustering [20]. A classical multidimensional scaling (MDS) analysis was carried out to help visualize the spatial locations of the clusters [21]. For a given positive integer k, the MDS algorithm determines a k-dimensional representation of the data space such that the distances between pairs of data points in the original data space are preserved as best as possible. We used k = 3 in our analysis to facilitate visualization. We used the OneClassPlusSVM.R script. This script is available at ftp://ftp-trace.ncbi.nih.gov/giab/ftp/technical/svclassify_Manuscript/Supplementary_Information/svclassify/.

### One-class classification using L_1_ distance

The set of 4000 random sites representing the class of likely non-SVs with a size range of 50 bps to 997527 bps were used for training the one-class classifier. First, a separate classifier was developed using data from each sequencing technology for these 4000 sites. The classifier was based on the empirical distribution of L_1_ distances of each of the 4000 sites from the mean M for the 4000 sites. For these likely non-SVs, a threshold value tp was determined such that a proportion ρ of the 4000 L_1_ distances were less than or equal to tp. The region R is then defined as the set of all points in the transformed data space whose L_1_ distance from the mean M is less than or equal to tp. When there are only two annotations measured for each site, this region takes the shape of a rhombus. In the high dimensional data space the shape of the region R is a multidimensional rhombus. The classification rule is as follows. Given any new site, calculate its L_1_ distance from M. If it is greater than tp classify it as a SV. Otherwise call it a non-SV. Five classifiers were developed one for each of the four sequencing technologies and one using the combined data. We used the Unsupervised.R script. This script is available at ftp://ftp-trace.ncbi.nih.gov/giab/ftp/technical/svclassify_Manuscript/Supplementary_Information/svclassify/.

### One-class classification using one-class SVM

Support Vector Machines (SVM) [22] are generally used for supervised learning when it is desired to develop a classification rule for classifying sites into two or more classes. Different versions of SVMs have been developed for one-class classification [23, 24]. We use the version proposed by Schölkopf *et al.* just as in the case of L_1_ one-class classification discussed above, we develop five classifiers based on data from each of the four sequencing technologies and a classifier based on the combined data from all four sequencing technologies to distinguish SVs from random regions and SVs from validated/assembled sets. In this analysis, a different data transform method was applied to each annotation. First, for each annotation we defined the deviation directions of interest compared to the reference distribution of SVs from the random regions to define outliers. According to the defined directions of deviations, we transformed the data so that the range of each annotation satisfies the required condition of one-class SVM. i.e. for each annotation, the larger the directional deviation was, the closer to 0 the transformed value was. One-class SVM implemented with e1071 package of the Comprehensive R Archive Network was trained by the transformed data of 4000 random regions to define linear class boundaries that may discriminate true SVs from randomly generated SVs. The proportion of SVs in the training set identified as outliers (false positive rate) 1-p was approximately controlled by a factor v in the training algorithm defined by the authors (Supplementary information 1). We used the OneClassPlusSVM.R script. This script is available at ftp://ftp-trace.ncbi.nih.gov/giab/ftp/technical/svclassify_Manuscript/Supplementary_Information/svclassify/.

### Ensemble classifiers

Above, an L_1_ classifier was developed separately for data from each of the four sequencing datasets. A fifth classifier was developed by combining annotations from all four datasets into a single model. Rather than combining the datasets, we can combine the four classifiers using an idea referred to as ensemble learning. We consider ensemble classifiers that are based on declaring a new site to be a SV provided at least *k* of the individual classifiers predict the site as a SV. We can do this for *k*=1, 2, 3, 4. These ensemble classifiers arising from the four L_1_ classifiers were investigated and their performances are reported in the results section. A similar process was repeated for the one-class SVM classifier. As the class boundaries developed with one-class SVM could have intersections, in one-class SVM analysis, for each SV, we recorded the smallest true negative rate of the training that lead to a classifier defines this SV as one from the random regions, as an equivalent to the proportion ρ used for the L_1_ classifiers.

We chose k=3 from the L_1_ classifier to produce our final high-confidence SVs, since we expect classifications based on evidence from multiple datasets are more likely to be robust. Candidate SV sites from Personalis, 1000 Genomes, and Spiral Genetics as well as Random Genome sites were stratified into sites with varying levels of evidence for an SV using the L_1_ classifier. To exclude difficult regions in which our classifier may give misleading results, we first excluded sites with Platinum Genomes coverage > 300 in the left and right flanking regions (~1.5 times the mean coverage, so these may be inside duplicated regions), as well as sites with Platinum Genomes mean mapping quality < 30 in the left or right flanking regions. We used the OneClassPlusSVM.R script. This script is available at ftp://ftp-trace.ncbi.nih.gov/giab/ftp/technical/svclassify_Manuscript/Supplementary_Information/svclassify/.

### Manual inspection of SVs

To understand the accuracy of our classifier, we manually inspected a subset of the sites from each call set. Specifically, we inspected all 17 random sites with ρ > 0.99 to determine if these might be real SVs. We also randomly selected 20 sites each from Personalis and 1000 Genomes with ρ > 0.99, and 10 sites from Personalis and 1000 Genomes with ρ < 0.68, 0.68 < ρ < 0.90, and 0.90 < ρ < 0.99 (or we inspected all sites if there were fewer than 10 in any category). Manual inspection was performed using the GeT-RM project browser (http://www.ncbi.nlm.nih.gov/variation/tools/get-rm/browse/), the integrative genomics viewer (IGV) (version 2.3.23 (26)) [25] and svviz (version 1.0.9; https://github.com/svviz/svviz) [10]. We selected the following tracks on GeT-RM Browser for manual inspections: GRCh37.p13 (GCF_000001405.25) Alternate Loci and Patch Alignments, GRC Curation Issues mapped to GRCh37.p13, Repeats identified by RepeatMasker, 1000 Genomes Phase 1 Strict Accessibility Mask, dbVar ClinVar Large Variations, dbVar 1000 Genomes Consortium Phase 3 (estd214), NIST-GIAB v.2.18 abnormal allele balance, NIST-GIAB v.2.18 calls with low mapping quality or high coverage, NIST-GIAB v.2.18 evidence of systematic sequencing errors, NIST-GIAB v.2.18 local alignment problems, NIST-GIAB v.2.18 low coverage, NIST-GIAB v.2.18 no call from HaplotypeCaller, NIST-GIAB v.2.18 regions likely have paralogs in the 1000 Genomes decoy, NIST-GIAB v.2.18 regions with structural variants in dbVar for NA12878, NIST-GIAB v.2.18 Simple Repeats from RepeatMasker, NIST-GIAB v.2.18 support from < 3 datasets after arbitration, NIST-GIAB v.2.18 uncertain regions due to low coverage/mapping quality. We observed coverage of the regions, numbers of soft-clipped reads, numbers of reads with deletions relative to the reference genome and numbers of SNPs/indels in the regions from Moleculo and PacBio aligned bam files using IGV.

### svviz

svviz (version 1.0.9; https://github.com/svviz/svviz) was used to visualize all four whole-genome sequencing data sets to see if there is support for a given structural variant [10]. It uses a realignment process to identify reads supporting the reference allele, reads supporting the structural variant (or alternate allele), and reads that are not informative one way or the other (ambiguous). svviz batch mode was used with default parameters to calculate summary statistics for SVs and non-SVs. In addition, inserted sequences were included as an input for svviz for Spiral Genetics’ insertions calls. For PacBio sequencing data, svviz’s “pacbio” optional parameter was used to retain lower quality alignments as support for the reference and alternate alleles since PacBio sequencing has a relatively high error rate. svviz’s commands, input files and output files are provided in svviz.zip.

### Trio and curated call sets

Read sequences for the trio analysis were downloaded from the Illumina Platinum Genomes project (http://www.illumina.com/platinumgenomes/) deposited at the European Nucleotide Archive (http://www.ebi.ac.uk/ena/data/view/ERP001960). The study accession was PRJEB3381 and the sample accessions were SAMEA1573618 for NA12878, SAMEA1573615 for NA12891, and SAMEA1573616 for NA12892. Sequences were aligned with BWA-MEM version 0.7.12. Deletion calls were reported by MetaSV [5] version 0.3 integrating Pindel [26] version 0.2.5a8, CNVnator [27] version 0.3, BreakDancer [28] version 1.4.5, and BreakSeq [29, 30] version 2.0-alpha. The curated, validated call set was generated from the 1000 Genomes Project [12], resulting in a set of 2406 calls which were previously reported for NA12878.

### 30x coverage dataset

We have performed down-sampling of the Platinum Genomes BAM file to 30x (which is 15% of the 200x data) by using samtools (version 0.1.19-44428cd) (samtools view -s 0.15) which reflects a typical sequencing project (e.g. Illumina HiSeq, 100 bp paired end reads with 30x coverage).

## Availability of supporting data

The data supporting the results of this article can be downloaded at ftp://ftp-trace.ncbi.nih.gov/giab/ftp/technical/svclassify_Manuscript/Supplementary_Information/.

## Authors’ contributions

HP, MP, GB, WL, JMZ, and MS designed the study.

HP wrote and performed the svclassify annotation script.

HP, MM, HYKL, HI, DC, GB, and JMZ wrote the manuscript.

MM and HYKL performed the MetaSV confirmation analyses.

MP and GB generated the high-confidence calls and PCR validation from Personalis.

NS, HP, and JMZ designed and performed the svviz analysis.

HI, DC, and WL performed the machine learning analysis.

All authors read and approved the final manuscript.

## Additional files

**Supplementary figure 1:**
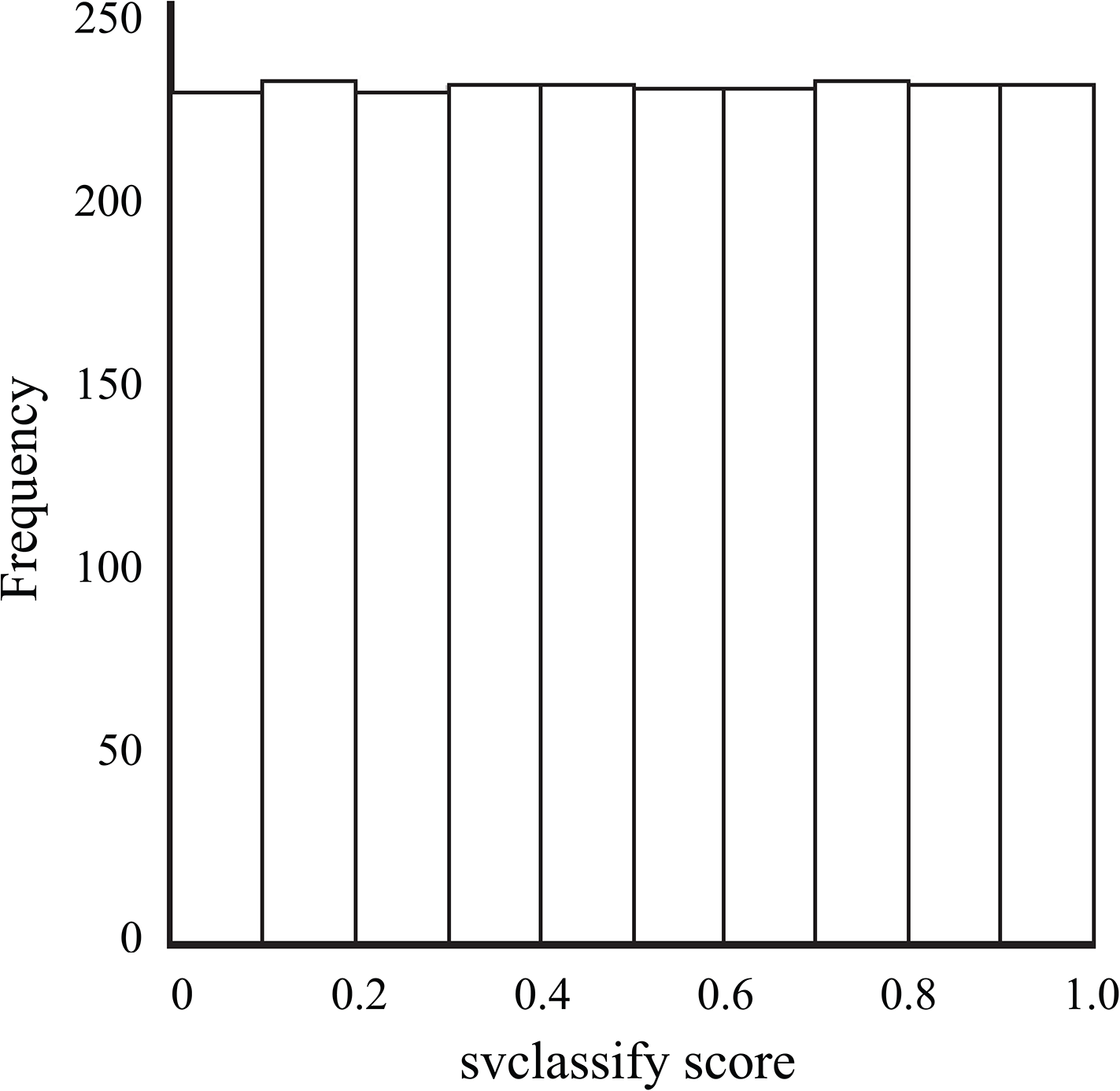
Histogram of ρ score for 2306 random regions with same size distribution as Personalis SVs.

**Supplementary figure 2:**
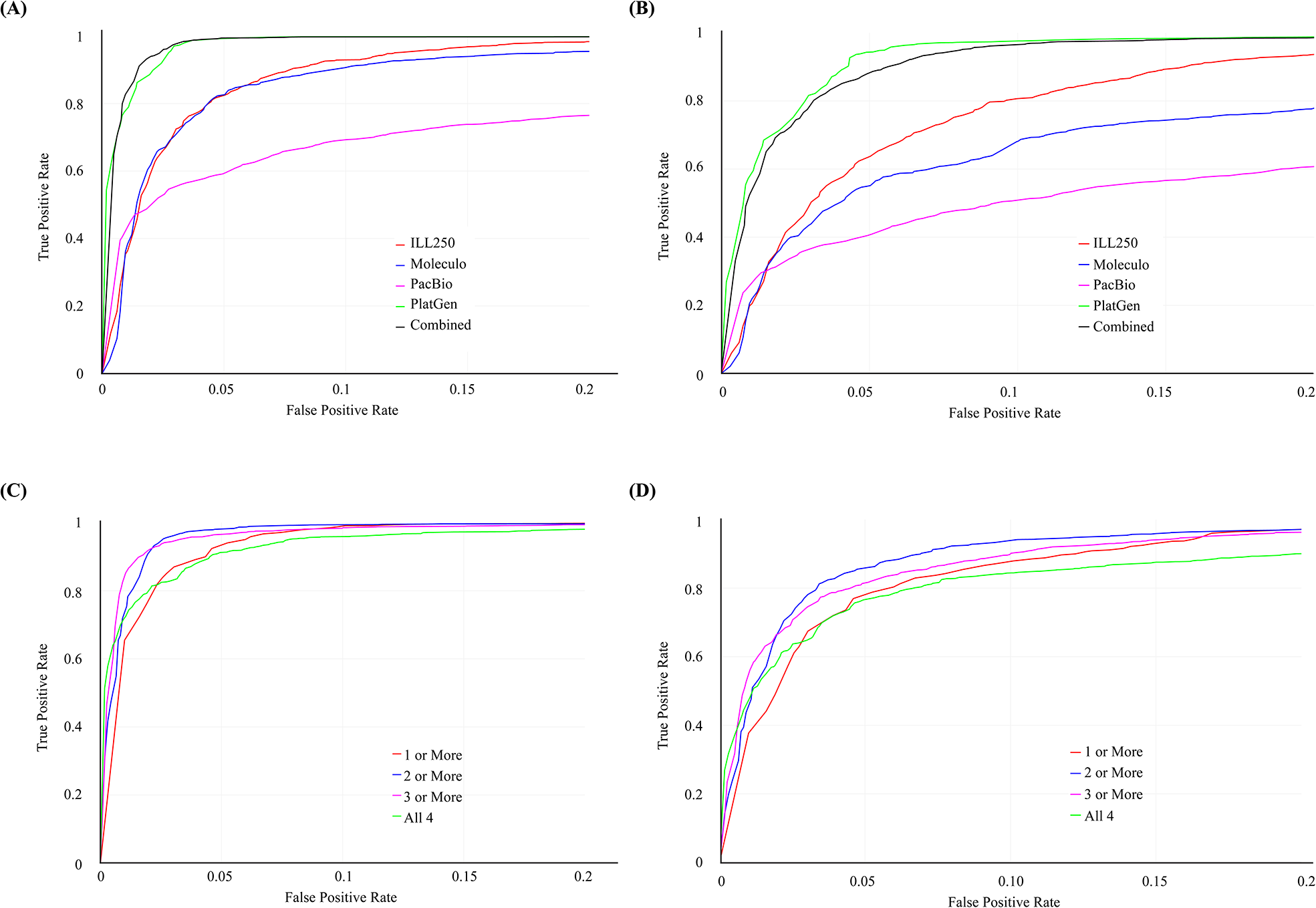
ROC curves for One-class classification using SVM, treating the 4000 Random regions as negatives and the Personalis or 1000 Genomes calls as positives. (A) ROC curves for one-class models for each dataset separately and for all combined for the Personalis validated deletion calls. (B) ROC curves for one-class models for each dataset separately and for all combined for the 1000 Genomes validated deletion calls. (C) ROC curves for one-class model requiring 1 or more, 2 or more, 3 or more, or all 4 technologies to have high classification scores for the Personalis validated deletion calls. (D) ROC curves for one-class model requiring 1 or more, 2 or more, 3 or more, or all 4 technologies to have high classification scores for the 1000 Genomes validated deletion calls. See original data at https://plot.ly/233/~parikhhm/, https://plot.ly/242/~parikhhm/, https://plot.ly/250/~parikhhm/, and https://plot.ly/258/~parikhhm/.

**Supplementary figure 3:**
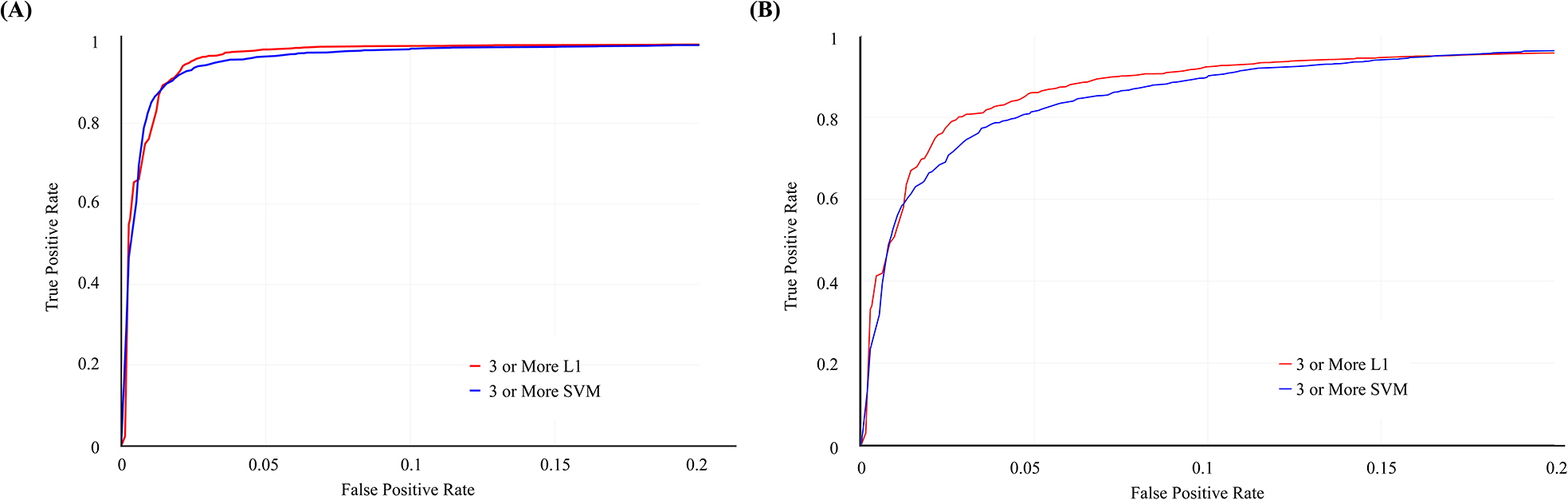
ROC curves for One-class classification using SVM and L_1_ “3 or more” strategy, treating the 4000 random regions as training negatives, treating (A) the Personalis deletion calls and (B) the 1000 Genomes deletion calls as testing positives and treating the 2306 random regions as testing negatives. See original data at https://plot.ly/266/~parikhhm/, and https://plot.ly/274/~parikhhm/.

**Supplementary figure 4:**
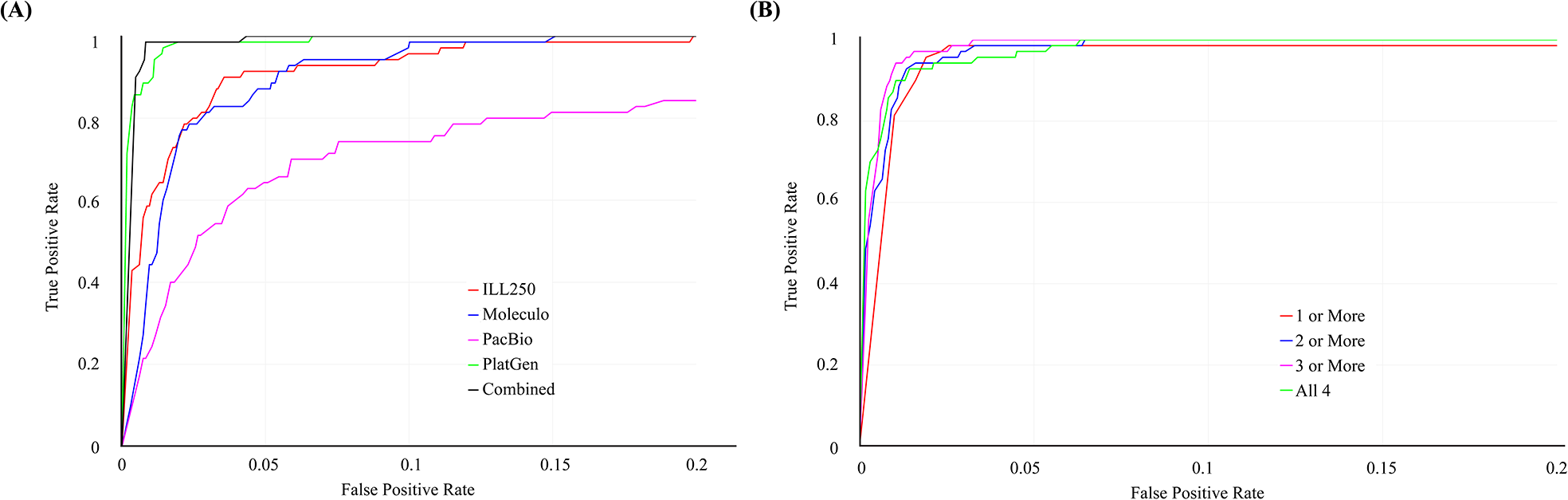
ROC curves for One-class classification using SVM, treating the 4000 random regions as negatives and the Spiral Genetics insertions calls as positives. (A) ROC curves for one-class models for each dataset separately and for all combined. (B) ROC curves for one-class model requiring 1 or more, 2 or more, 3 or more, or all 4 technologies to have high classification scores. See original data at https://plot.ly/~desuchen0929/391, and https://plot.ly/297/~parikhhm/.

**Supplementary figure 5:**
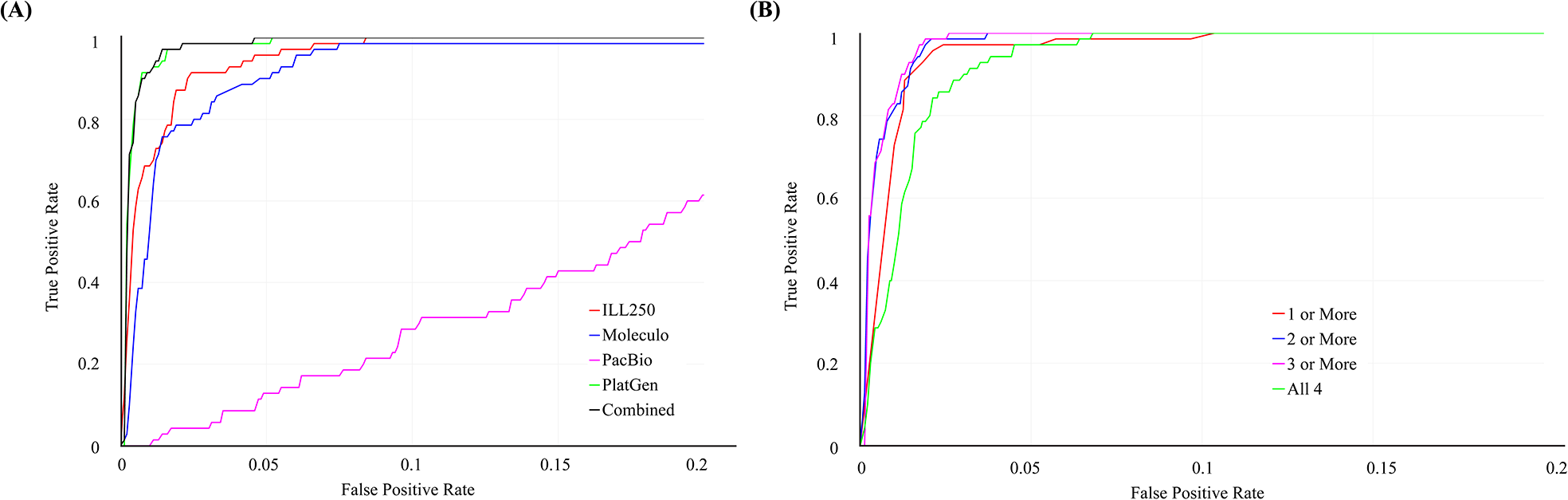
ROC curves for One-class classification using the L_1_ Distance, treating the 4000 random regions as negatives and the Spiral Genetics insertions calls as positives. (A) ROC curves for one-class models for each dataset separately and for all combined. (B) ROC curves for one-class model requiring 1 or more, 2 or more, 3 or more, or all 4 technologies to have high classification scores. See original data at https://plot.ly/~desuchen0929/386, and https://plot.ly/330/~parikhhm/.

**Supplementary figure 6:**
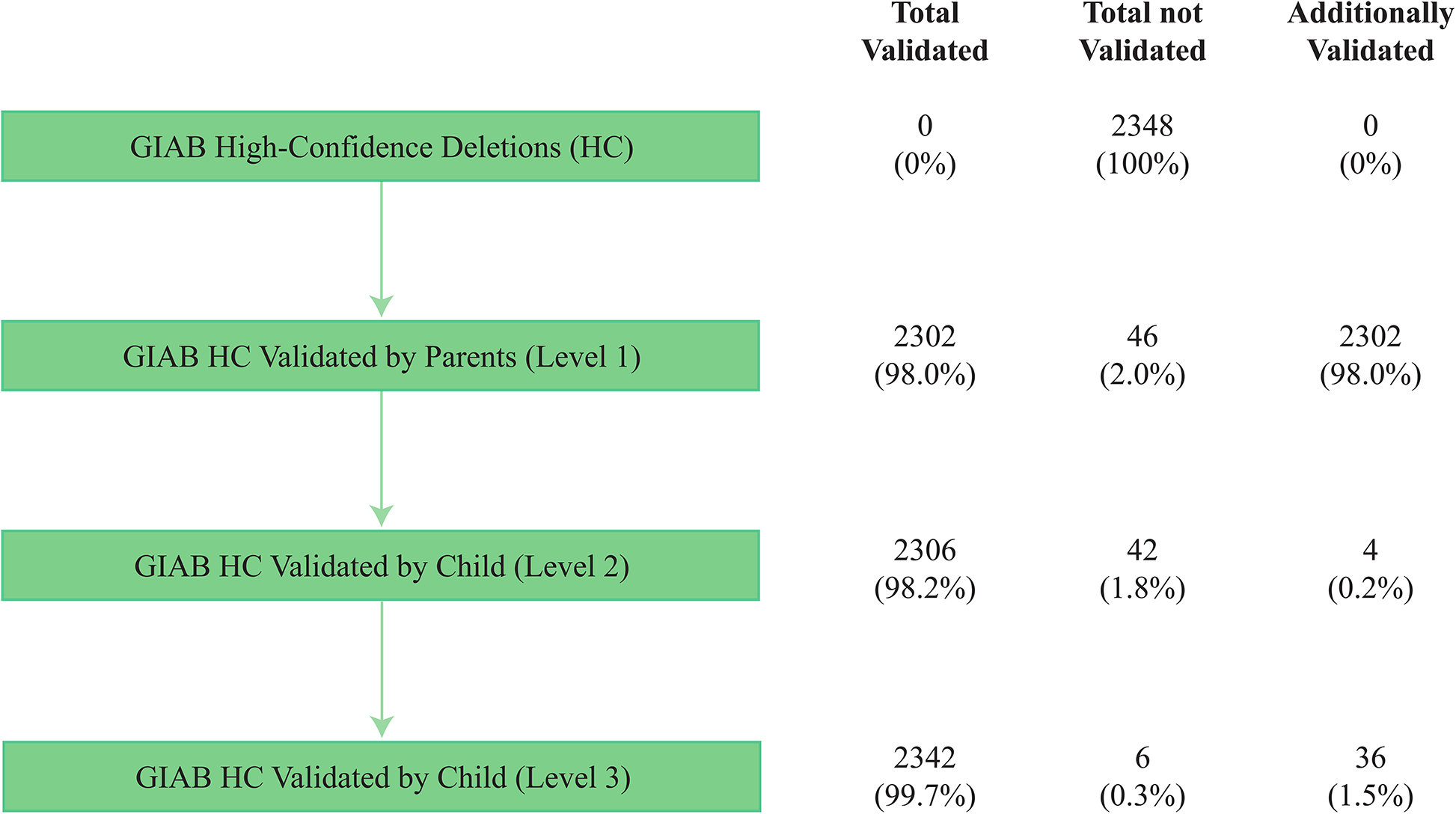
Three levels of validation with trio analysis. “Total Validated” gives the number of sites validated by that level or any level above. “Total not Validated” gives the number of sites not validated in that level or any level above. “Additionally Validated” gives the number of sites validated by that level, excluding sites validated in levels above.

**Supplementary figure 7:**
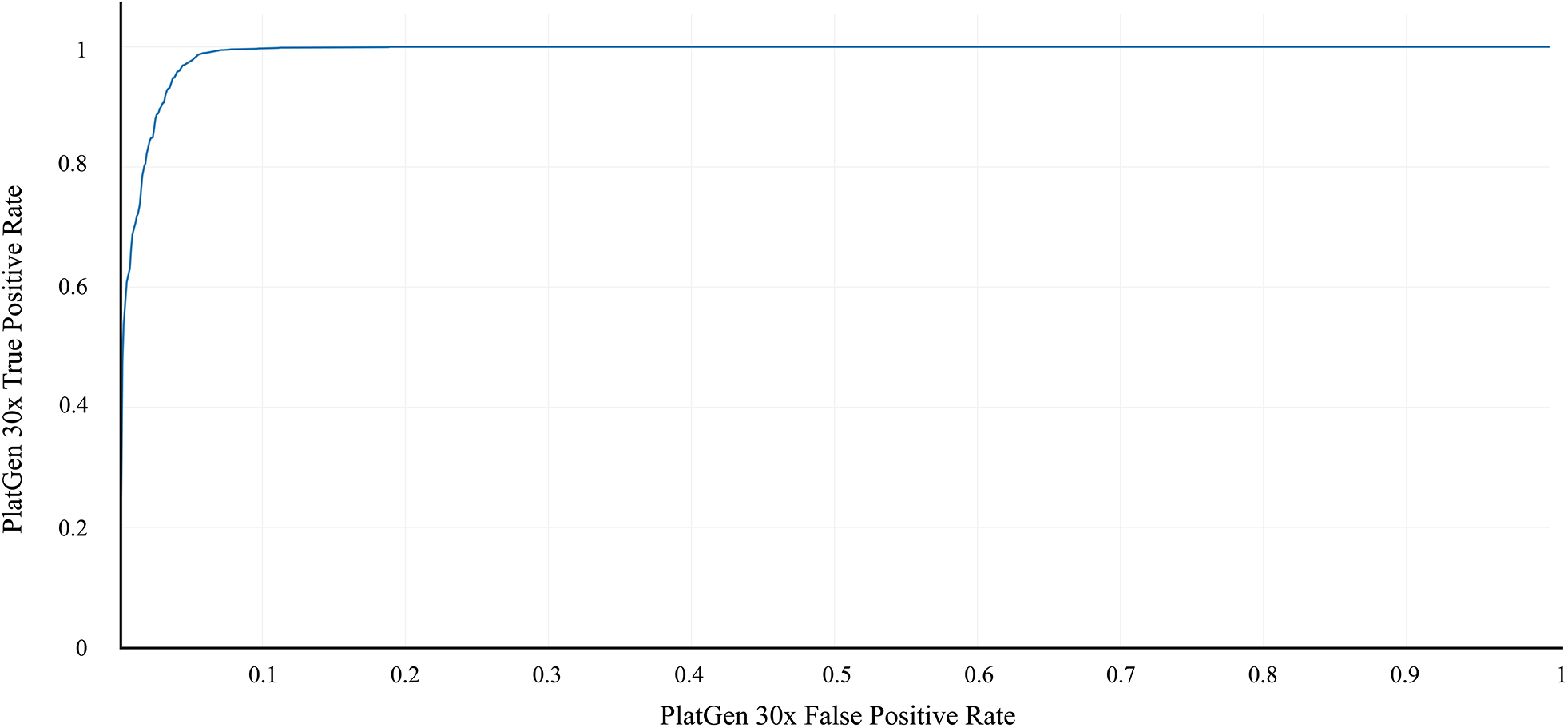
ROC curves for One-class classification using the L_1_ Distance, treating the 2306 Random regions (size distribution matching to Personalis) as negatives and the Personalis as positives using the Platinum down-sampling 30x data. See original data at https://plot.ly/165/~parikhhm/.

**Supplementary table 1:** Annotations for each of the SV calls as well as likely non-SV regions from the Platinum Genomes 2x100bps HiSeq aligned sequence dataset for NA12878 using svclassify.

**Supplementary table 2:** Annotations for each of the SV calls as well as likely non-SV regions from the Illumina HiSeq (read length = 250 bps) aligned sequence dataset for NA12878 using svclassify.

**Supplementary table 3:** Annotations for each of the SV calls as well as likely non-SV regions from the Moleculo aligned sequence dataset for NA12878 using svclassify.

**Supplementary table 4:** Annotations for each of the SV calls as well as likely non-SV regions from the PacBio aligned sequence dataset for NA12878 using svclassify.

**Supplementary table 5:**
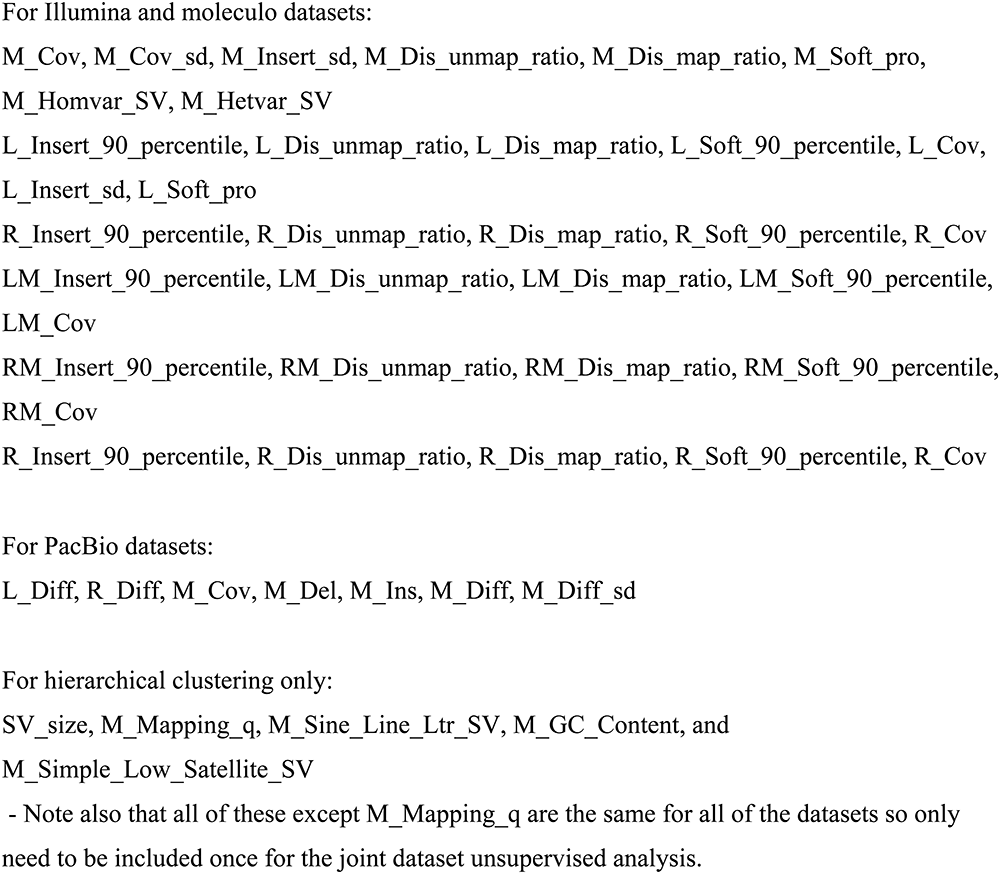
Selected characteristics for hierarchical clustering and One-class models.

**Supplementary table 6:**
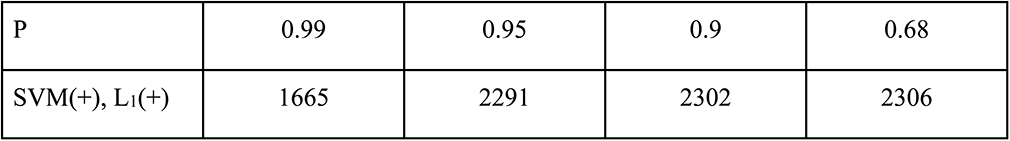
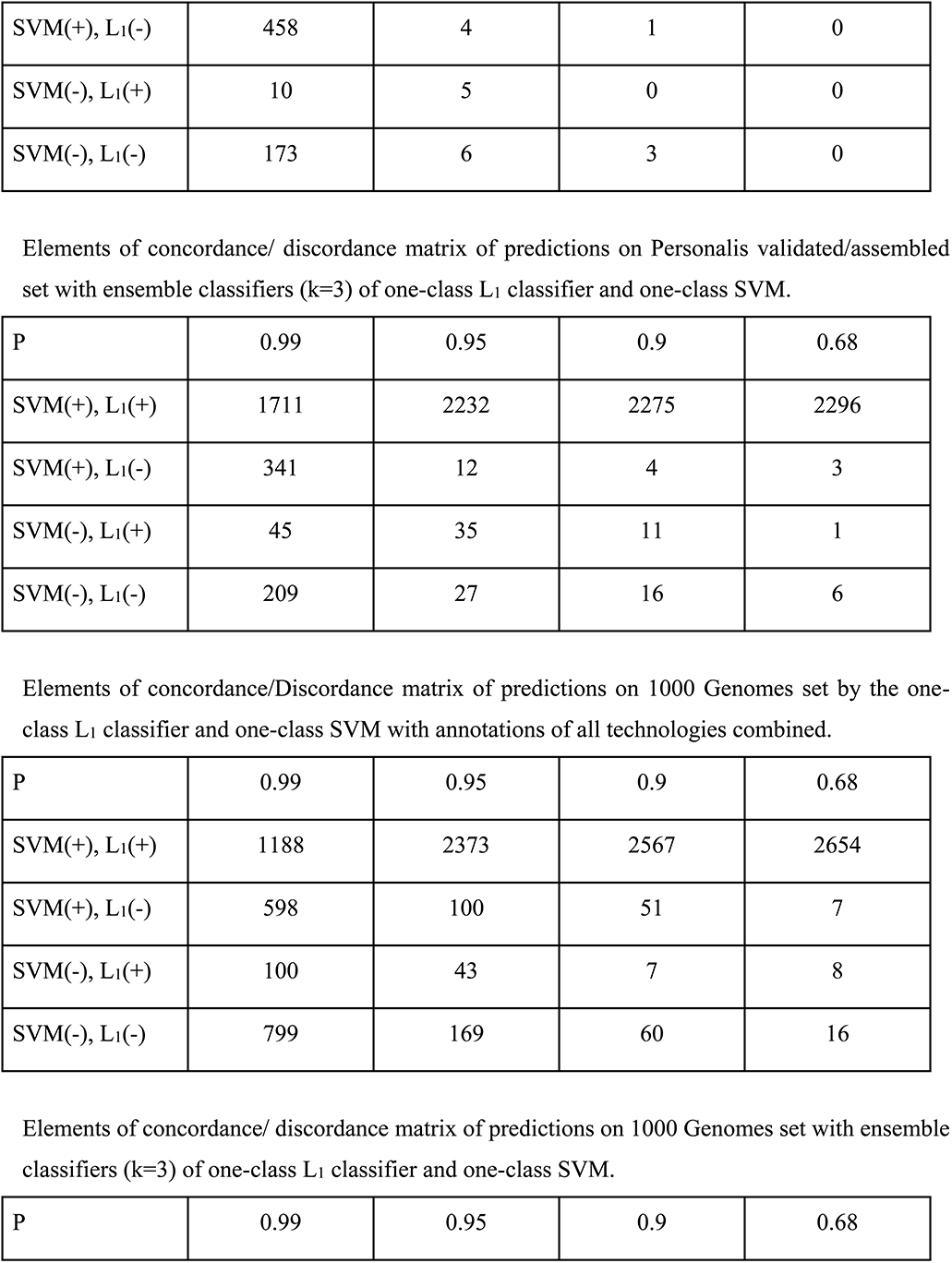
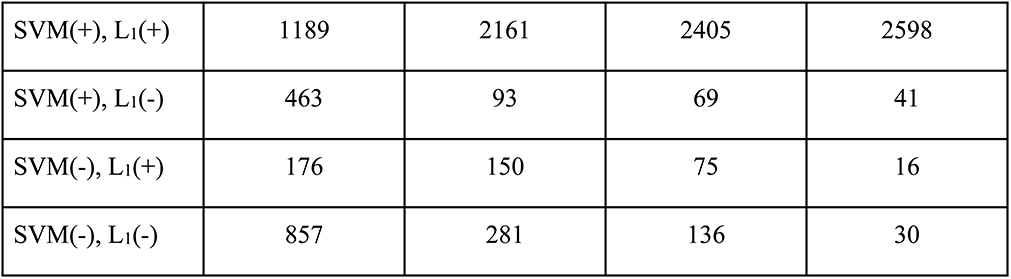
Elements of concordance/discordance matrix of predictions on Personalis validated/assembled set by the one-class L_1_ classifier and one-class SVM with annotations of all technologies combined.

|  |  |  |  |  |
| --- | --- | --- | --- | --- |
| P | 0.99 | 0.95 | 0.9 | 0.68 |
| SVM(+), $L_1$(+) | 1665 | 2291 | 2302 | 2306 |
| SVM(+), L<sub>1</sub>(-) | 458 | 4 | 1 | 0 |
| SVM(-), L<sub>1</sub>(+) | 10 | 5 | 0 | 0 |
| SVM(-), L<sub>1</sub>(-) | 173 | 6 | 3 | 0 |
| P | 0.99 | 0.95 | 0.9 | 0.68 |
| SVM(+), L<sub>1</sub>(+) | 1711 | 2232 | 2275 | 2296 |
| SVM(+), L<sub>1</sub>(-) | 341 | 12 | 4 | 3 |
| SVM(-), L<sub>1</sub>(+) | 45 | 35 | 11 | 1 |
| SVM(-), L<sub>1</sub>(-) | 209 | 27 | 16 | 6 |
| P | 0.99 | 0.95 | 0.9 | 0.68 |
| SVM(+), L<sub>1</sub>(+) | 1188 | 2373 | 2567 | 2654 |
| SVM(+), L<sub>1</sub>(-) | 598 | 100 | 51 | 7 |
| SVM(-), L<sub>1</sub>(+) | 100 | 43 | 7 | 8 |
| SVM(-), L<sub>1</sub>(-) | 799 | 169 | 60 | 16 |
| P | 0.99 | 0.95 | 0.9 | 0.68 |
| SVM(+), L<sub>1</sub>(+) | 1189 | 2161 | 2405 | 2598 |
| SVM(+), L<sub>1</sub>(-) | 463 | 93 | 69 | 41 |
| SVM(-), L<sub>1</sub>(+) | 176 | 150 | 75 | 16 |
| SVM(-), L<sub>1</sub>(-) | 857 | 281 | 136 | 30 |

**Supplementary table 7:** Manual inspection characteristics of the 20 randomly selected Personalis sites with ρ > 0.99 of one-class results.

**Supplementary table 8:** Manual inspection characteristics of all 10 of the randomly selected Personalis sites with 0.9 < ρ < 0.99 of one-class results.

**Supplementary table 9:** Manual inspection characteristics of all 8 of the randomly selected Personalis sites with ρ < 0.9 of one-class results.

**Supplementary table 10:** Manual inspection characteristics of the 20 randomly selected 1000 Genomes sites with ρ > 0.99 of one-class results.

**Supplementary table 11:** Manual inspection characteristics of the 9 randomly selected 1000 Genomes sites with 0.9 < ρ < 0.99 of one-class results.

**Supplementary table 12:** Manual inspection characteristics of the 8 randomly selected 1000 Genomes sites with 0.7 < ρ < 0.9 of one-class results.

**Supplementary table 13:** Manual inspection characteristics of the 7 randomly selected 1000 Genomes sites with ρ < 0.7 of one-class results.

**Supplementary table 14:** Manual inspection characteristics of the insertions from Spiral Genetics using svviz.

**Supplementary table 15:** Number of overlapping deletion calls between Personalis and 1000 Genomes deletion calls with different amounts of overlap.

| **Personalis unique deletion calls** | **1000 Genomes unique deletion calls** | **Overlap** | **# of overlapping deletion calls** |
| --- | --- | --- | --- |
| 2336 | 1825 | 1 bp | 1082 |
| 2336 | 1825 | 10 % | 1082 |
| 2336 | 1825 | 25 % | 1081 |
| 2336 | 1825 | 50 % | 1076 |
| 2336 | 1825 | 75 % | 1070 |
| 2336 | 1825 | 90 % | 1066 |
| 2336 | 1825 | 100 % | 986 |

**Supplementary table 16:**
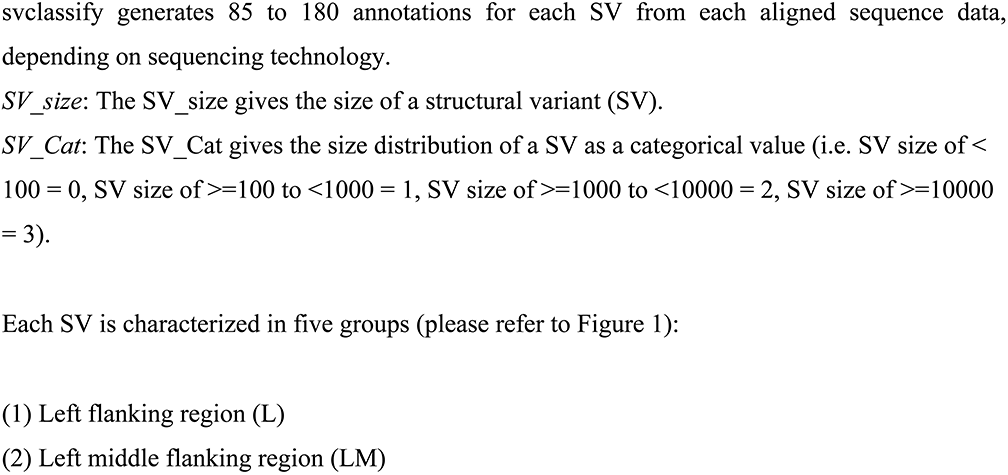
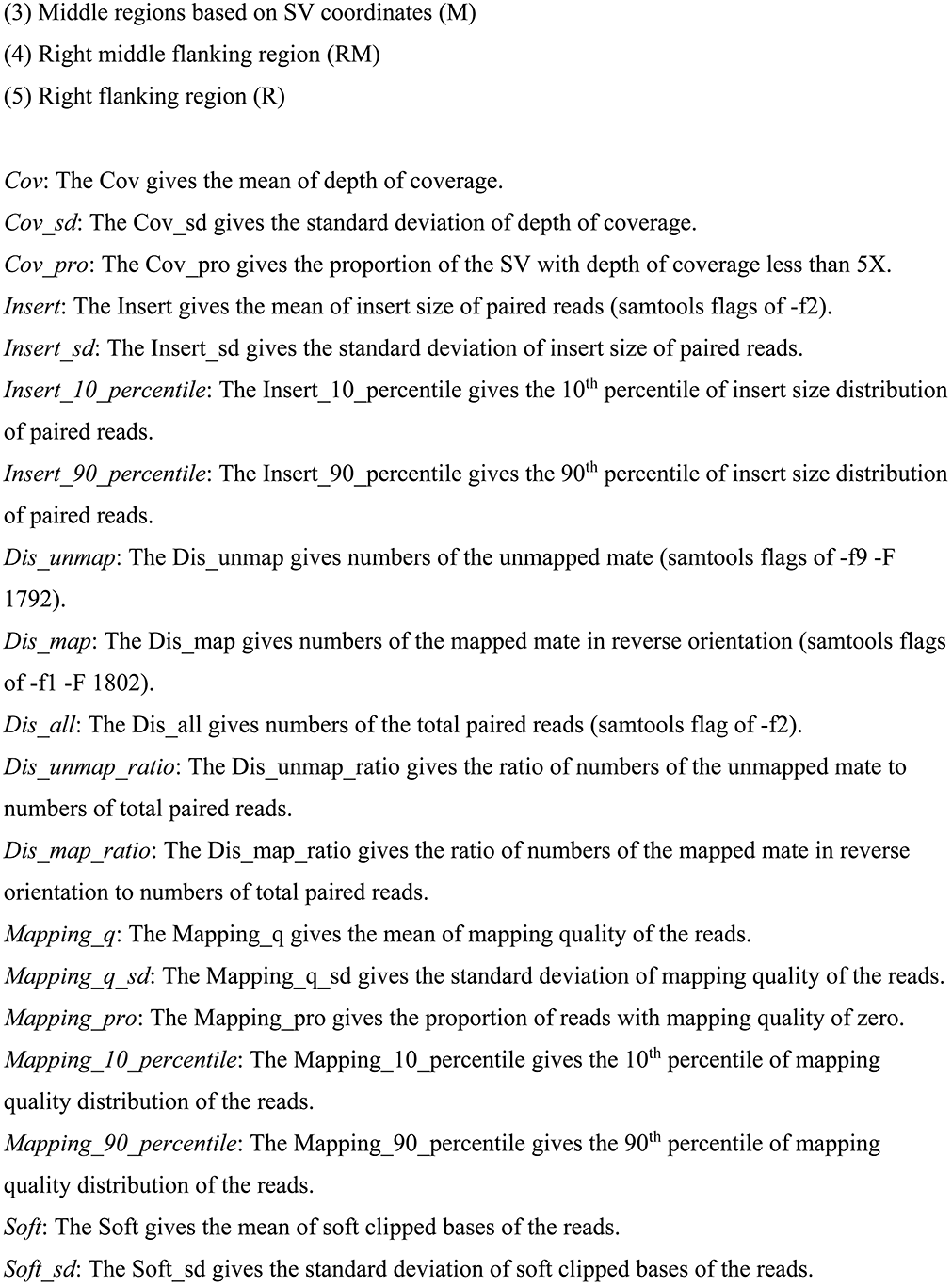
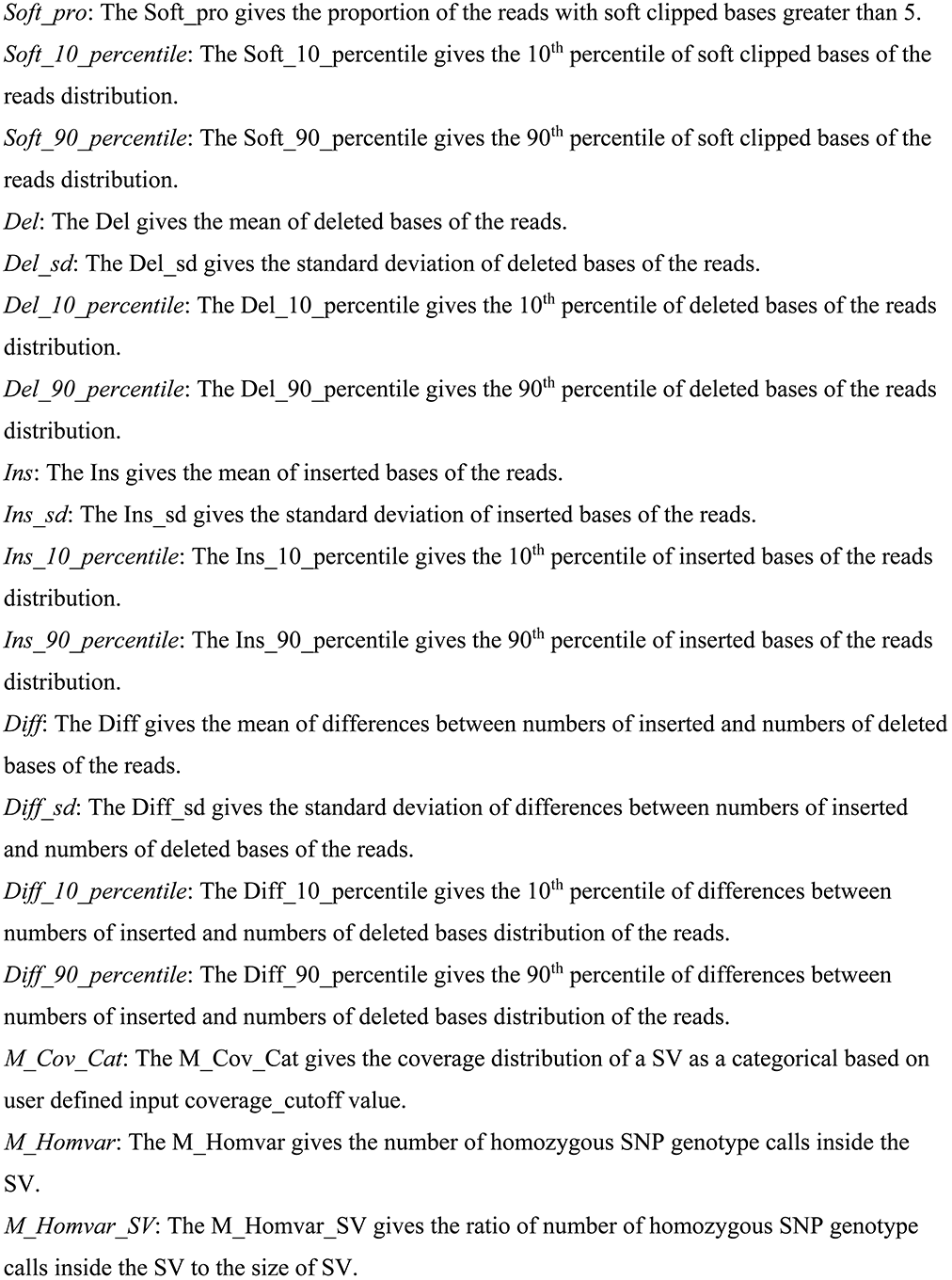
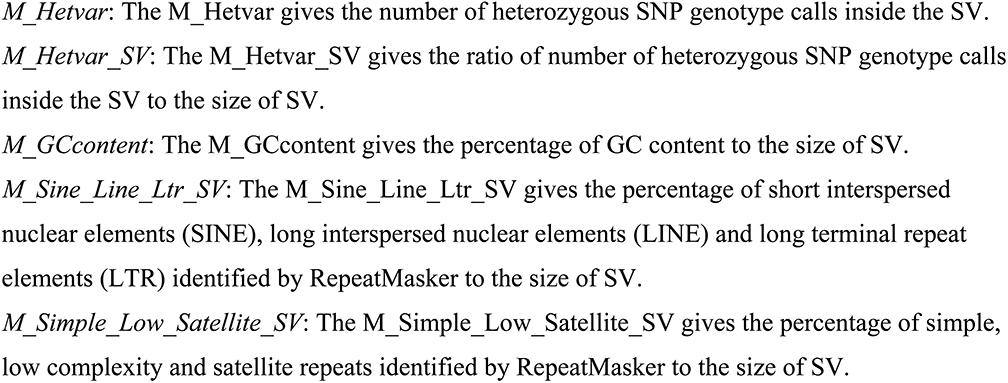
Output format of svclassify.

## Acknowledgements

The authors would like to thank Niranjan Shekar from Spiral Genetics for contributing the insertion calls used in this manuscript, and Jian Li from Bina Technologies for helping to compile the validated calls from 1000 Genomes. Certain commercial equipment, instruments, or materials are identified in this document and that such identification does not imply recommendation or endorsement by the National Institute of Standards and Technology, nor does it imply that the products identified are necessarily the best available for the purpose.

## Supplementary Information

### 1 Data transform for one-class SVM

For a certain annotation, the “right-tail” case means outliers should have positive deviations, the “left-tail” case means outliers should have negative deviations, and the “both-tail” case means that outliers could have either positive or negative deviations. Reference deviations were then calculated for different cases. For the left-tail and right-tail cases,

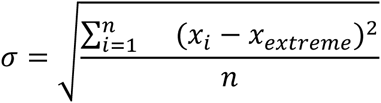

σ is the reference deviation, *x_i_* is annotation value of the *i*th site from the random regions. *n* is the number of sites from the random regions (*n* = 4000). *X_extreme_* is either the minimum of *x_i_* for the right-tail case, or the maximum of *x_i_* for left-tail case. For any observation *x* of the same annotation for any SV, the transform *y* is

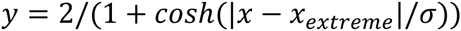

For both-tail case, we define two reference deviations σ_right_ and σ_left_ for either positive or negative deviations from the median *x_med_* of *x_i_,*

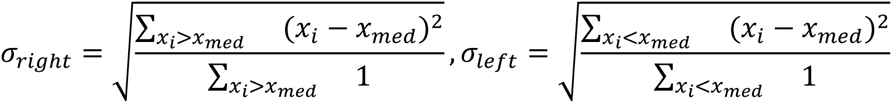

The transform *y* is

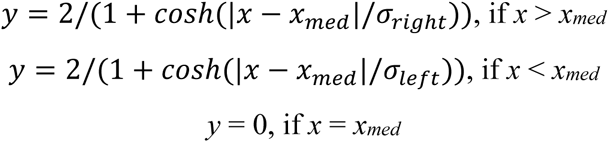

Therefore outliers indicating potential SVs approach 0 in this transform, which is required by the application of one-class SVM. In the transformed metric space, linear classifiers were trained by the one-class SVM (implemented with package e1071 in the Comprehensive R Archive Network) with SVs from the random regions as the training set. The proportion of SVs in the training set identified as outliers (false positive rate) *1-p* was approximately controlled by a factor v in the training algorithm defined by the authors. In short, *v* ∈ (0,1) defines the ratio of penalty induced by margin size (e.g. distance from origin point to the class boundary with linear kernel) and penalty induced by number of outliers in the training set in the total penalty function for soft margin case. Higher v allows more training data points to be on the outliers side to maximize the margin. Classifiers at different v’s were then applied to predict other SV data sets.

